# Antigen identification and high-throughput interaction mapping by reprogramming viral entry

**DOI:** 10.1101/2021.09.18.460796

**Authors:** Connor S. Dobson, Anna N. Reich, Stephanie Gaglione, Blake E. Smith, Ellen J. Kim, Jiayi Dong, Larance Ronsard, Vintus Okonkwo, Daniel Lingwood, Michael Dougan, Stephanie K. Dougan, Michael E. Birnbaum

## Abstract

Deciphering immune recognition is critical for understanding a broad range of diseases and for the development of effective vaccines and immunotherapies. Efforts to do so are limited by a lack of technologies capable of simultaneously capturing the complexity of adaptive immune receptor repertoires and the landscape of potential antigens. To address this, we present RAPTR (**R**eceptor-**A**ntigen **P**airing by **T**argeted **R**etroviruses). RAPTR combines viral pseudotyping and molecular engineering approaches to enable one-pot library on library interaction screens by displaying antigens on the surface of lentiviruses and encoding their identity in the viral genome. Antigen-specific viral infection of cells allows readout of both antigen and receptor identities via single-cell sequencing. The resulting system is modular, scalable, and compatible with any cell type, making it easily implemented. These techniques provide a suite of new tools for targeted viral entry, molecular engineering, and interaction screens with broad potential applications.

---

The adaptive immune system recognizes and responds to the diverse pathogens humans encounter throughout their lives. B cells recognize extracellular antigens via the B cell receptor (BCR), leading to the production of secreted antibodies to inhibit and eliminate pathogens. T cells use T cell receptors (TCRs) to recognize short peptide fragments presented by major histocompatibility complex proteins (pMHCs), enabling antigen-specific coordination of the immune response and elimination of malignant or infected cells. This remarkable ability to sense, respond to, and remember threats is key to any successful adaptive immune response, and accordingly is the cornerstone of successful vaccines and immunotherapies.

The importance of T cell-mediated immunity has led to the development of a series of approaches dedicated to identifying antigens or pMHC-TCR pairs^1^. Conventional T cell assays, such as ELISPOT or intracellular cytokine staining^2^, provide direct readouts of T cell function. While these approaches can be highly multiplexed, they do not readily lend themselves to TCR sequencing and are limited in their ability to identify single reactive antigens. More recently, T-Scan has broadened the antigenic scope of functional assays to genome scale^3^, but cannot provide paired receptor information without panning of pre-determined TCRs or iterated steps of target antigen identification followed by sorting of reactive cells for TCR sequencing. Other recent cell-based reporter assays convert pMHCs into the recognition domain of immune signaling complexes^4,5^. These approaches can identify successful TCR-pMHC interactions for single TCRs from antigen libraries on the scale of 10^3^-10^4^, but require substantial library redundancy, are limited in their ability to multiplex TCRs, and may require multiple rounds of screening. Requiring a discrete experiment per TCR imposes a significant scalability constraint; each individual has a TCR repertoire of ∼10^7^ unique clones^6,7^, and very little overlap is expected between individual repertoires even for people who share common MHC alleles.

Recombinant protein-based screening can broaden either the number of T cell clones or the number of antigens that are feasible to screen. Yeast display of pMHCs enables screening of ∼10^8^ unique pMHC antigens^8,9^, but can only examine a limited number of recombinantly expressed TCRs at a time, and can require significant optimization for each MHC allele. Conversely, recent advances in barcoded pMHC multimers enable screens on the order of 10^3^ antigens in bulk^10,11^, or hundreds while maintaining receptor-antigen pairing^12,13^. While such analyses can be performed on polyclonal T cells, they are inherently bottlenecked by several technical limitations: (i) the need to manually assemble individual barcoded multimers; (ii) the ability to correctly identify interactions in large pools of multimers; and (iii) the relatively small set of MHC molecules that have been recombinantly expressed successfully.

Identifying antigenic targets of B cells poses similar limitations. Recent approaches have shown that antigen-BCR pairs can be identified via the oligonucleotide tagging of recombinantly expressed proteins^14,15^, which is inherently limited in scale. Thus, our understanding of antigen recognition by both arms of the adaptive immune system is currently constrained by significant hurdles to experimental scale.

Deciphering the full complexity of immune recognition requires the ability to screen for interaction pairs while incorporating diversity of both antigen receptors and their targets at the same time. This represents a broader experimental challenge for interaction screens commonly known as “library vs library screening.” The most well-established approach to this problem is yeast two-hybrid^16^, in which intracellular protein pairs are used to drive expression of a reporter gene. Other previously established approaches include automated individual ELISA screens of recombinantly expressed protein pairs^17–19^, as well as mass-spectrometry-based identification of interacting pairs^20,21^. There have been several recent efforts at designing novel systems, including those based on yeast mating or spatial colocalization of DNA-barcoded molecules^22,23^. However, these approaches can be labor-intensive, are often not suited for complex, extracellular protein complexes such as immune receptors, and may be inefficient at low (i.e. micromolar) affinities. Together, these limitations have thus far rendered such approaches unsuitable for applications such as identifying TCR-pMHC pairs.

To overcome the above limitations, we have developed a technique that combines lentiviral surface display^24–26^ with a versatile pseudotyping strategy and viral genome engineering to enable one-pot library vs library screening (**Figure 1**). We demonstrate that our pseudotyping strategy – an engineered fusogen termed VSVGmut co-expressed with a targeting moiety – is general and versatile for both receptor and ligand usage. We leverage these abilities to present Receptor-Antigen Pairing by Targeted Retroviruses (RAPTR), which matches receptors with their cognate antigens based on specific infection of receptor-expressing cells by antigen-displaying viruses. Putative hits can be read out by bulk or single-cell sequencing, enabling screens of single or polyclonal receptors. We demonstrate the feasibility of this approach for both T and B cell receptors, including a library on library screen comprising 96 pMHC antigens and receptors enriched from a library of >450,000 TCRs (thousands of potential interactions).

**Figure 1.**
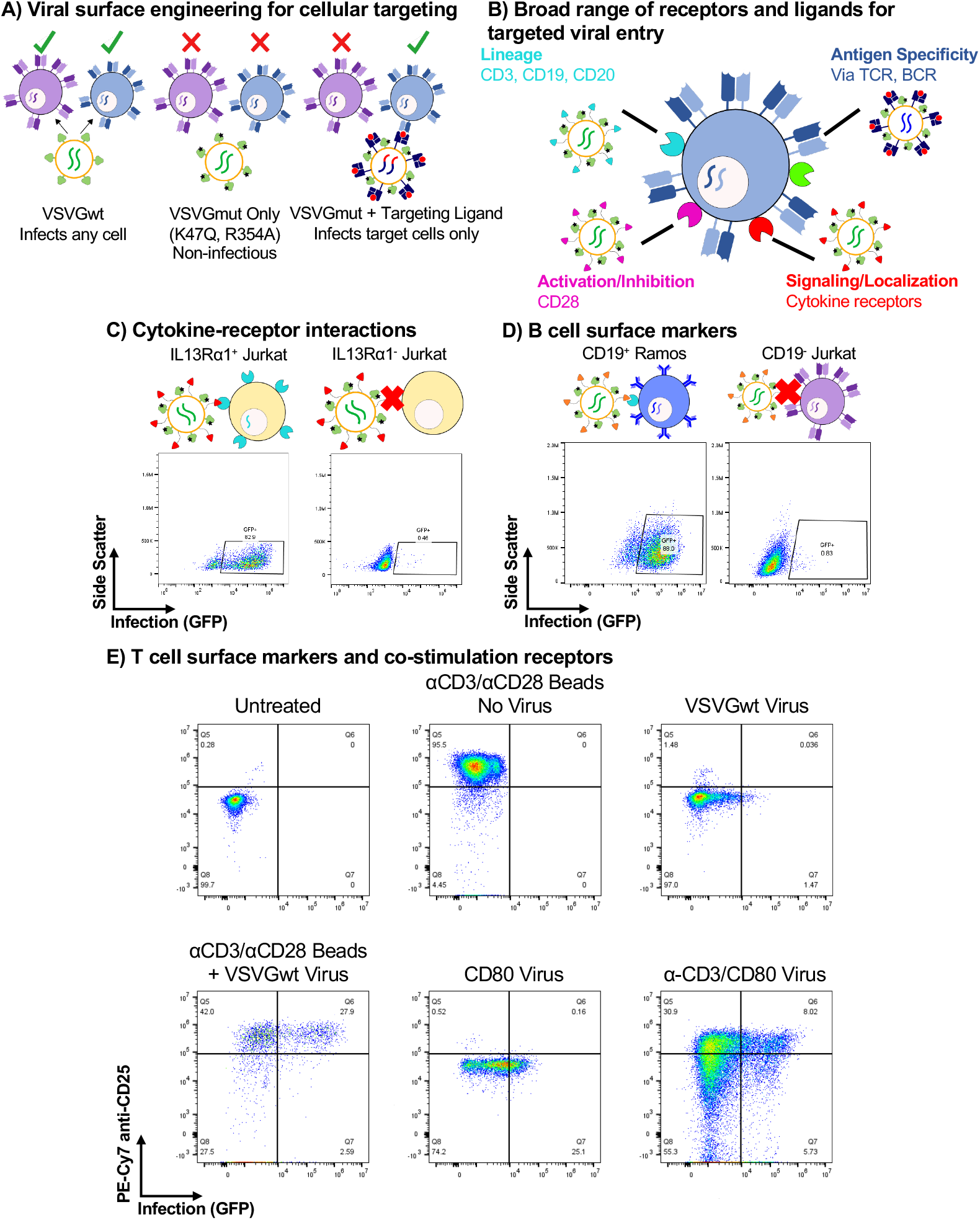
The VSVGmut pseudotyping system is modular and utilizes a broad range of receptors and ligands. **A**, VSVGmut (VSVG K47Q, R354A) pseudotyping system enables specific targeting by coexpression of receptor-blinded VSV-G and a modular targeting ligand. **B**, VSVGmut is compatible with a broad range of receptors and targeting ligands, including immune cell surface markers, signaling and antigen receptors. **C**, Specific infection of IL-13Rα1-expressing Jurkat cells (left) as compared to parental cell line (IL-13Rα1^-^, right) via VSVGmut lentiviruses displaying surface tethered IL-13; data representative of 2 biologically independent experiments. **D**, Specific infection of CD19^+^ Ramos cells (left), but not CD19^-^ Jurkat cells (right) by VSVGmut pseudotyped viruses displaying an anti-CD19 scFv; data are representative of 2 biologically independent experiments. **E**, Infection of Primary CD8 T cells (MOI = 1) via lentiviruses displaying anti-CD3 Fab and/or the costimulatory receptor CD80, or VSVGwt viruses with or without anti-CD3/anti-CD28 magnetic beads; data were collected at day 4 post-infection and are representative of three biological replicates.

## Results

### VSVGmut pseudotyped lentiviruses enable modular tropism

Developing a scalable pipeline for antigen-receptor screening presents several requirements: (i) a straightforward ability to track both receptor and antigen sequences; (ii) mammalian expression systems to maximize the ability to express complex, multimeric proteins; and (iii) the ability to generate selection reagents without the need to recombinantly express and characterize each protein for every experiment. To address these needs, we turned to lentiviruses, which have decades of precedent for molecular manipulations. They can be created at large scale, are already known to infect T cells, and, upon successful infection, leave a permanent record of infection due to their integration into the host genome.

Lentiviruses have been pseudotyped via a number of different strategies to enable their use as biotechnology tools and gene therapy vectors^27^. Due to its robustness and efficient infection of many cell types, vesicular stomatitis virus G protein (VSVG, referred to as VSVGwt in this manuscript) is the most common pseudotype for laboratory studies and cell manufacturing for clinical applications^28^. More recently, approaches to enable cell type-specific targeting, via co-expressing receptor-blinded versions of Sindbis virus^29– 31^ or paramyxovirus envelope proteins^32–37^ with targeting moieties have been described. While these approaches show promise, they have been reported to have strict limitations on targeting ligand and receptor binding topology due to their mechanism of entry.

Since these factors constrain the generality of an interaction screening system, we developed an alternative strategy based upon VSVG. We used the recently described crystal structure of VSVG in complex with its native receptor^38^, the low-density lipoprotein receptor (LDLR), to engineer VSVGmut, which incorporates the K47Q and R354A mutations reported to ablate affinity for the LDLR family of receptors (**Figure 1A**). To retarget VSVGmut pseudotyped viruses, we co-expressed a variety of surface-bound molecules during viral production (**Figure 1B**). To benchmark against established systems^33^, we used IL-13 as a viral targeting ligand. After testing several surface architectures (**Supplementary Figure 1**), we found that while many distinct constructs conferred specific infection of IL-13Rα1-expressing cells, the dimerized, surface-tethered IL-13 yielded the most efficient infection while retaining excellent specificity (**Figure 1C**). We also found that VSVGmut pseudotyped viruses displaying an anti-CD19 single-chain antibody fragment (scFv) efficiently infected CD19^+^ B cell lines, but not CD19^-^ T cell lines (**Figure 1D**).

Next, we sought to exploit the unique modularity of the VSVGmut system to develop combinatorial targeting strategies. While display of the anti-CD3 Fab UCHT1 yielded only modest infection of Jurkat T cell lines, CD80 mediated robust infection (**Supplementary Figure 2A**). Although the infection rate for the combinatorial strategy was not better than for CD80 alone, we reasoned that the ability to incorporate multiple signals during infection could be useful for engineering primary cells by providing user-defined phenotypic inputs. Therefore, we used these viruses to infect primary CD8 T cells and measured infection and CD25 upregulation as a marker of activation (**Figure 1E**). We observed that the anti-CD3/CD80 combination viruses both infected and activated cells, but viruses displaying CD80 alone infected but did not activate the cells. To our knowledge, this is the first example of a combinatorial viral targeting system reported to date.

### Antigen-specific T cell targeting

Following the observation that anti-CD3 viruses yielded infection of T cells via a component of the TCR complex, we next sought to determine whether we could achieve antigen-specific cell entry via the TCR itself. One potential challenge to repurposing the TCR-pMHC interaction for viral entry is its affinity, which is typically 1-50 μM^39^. We therefore generated cell lines expressing a range of previously characterized affinity variants for the 1G4 TCR^40,41^, which recognizes a peptide derived from the NY-ESO-1 cancer testis antigen (SLLMWITQV) presented in the context of HLA-A*02:01, with reported affinities ranging from picomolar to micromolar (**Figure 2A**). To display these molecules on the virus surface, we expressed them as single-chain trimers^42^, which are comprised of covalently linked peptide, beta-2-microglobulin, and MHC. We found that pMHC-displaying lentiviruses are able to efficiently infect T cells in an antigen-dependent manner. We observed similar infection efficiency across the tested affinity range, with only a modest reduction for the lowest-affinity variant (**Figure 2B**). To ensure that TCR-mediated infection was generalizable, we displayed several individual pMHCs as single-chain trimers alongside VSVGmut and used them to infect either Jurkat cells expressing its parental TCR (off-target) or on-target J76 cell lines expressing their cognate TCRs (**Figure 2C**). We observed specific infection across 3 different TCR-pMHC pairs with minimal background infection. As VSVG enables cell entry via an endocytic route^43^, and because endocytosis is one of the earliest downstream events following T cell activation^44–46^, we next sought to determine whether viral infection was accompanied by TCR signaling. We measured CD69 expression following viral infection (**Figure 2D**), and observed robust CD69 upregulation during TCR-mediated viral entry, but not during off-target combinations. Moreover, TCR-mediated entry was inhibited by dasatinib, a TCR signaling inhibitor^47^, while TCR-independent infection via VSVGwt was unaffected (**Supplementary Figure 2B**). Thus, we concluded that pMHC-targeted viruses integrate both binding and signaling as a means of infection.

**Figure 2.**
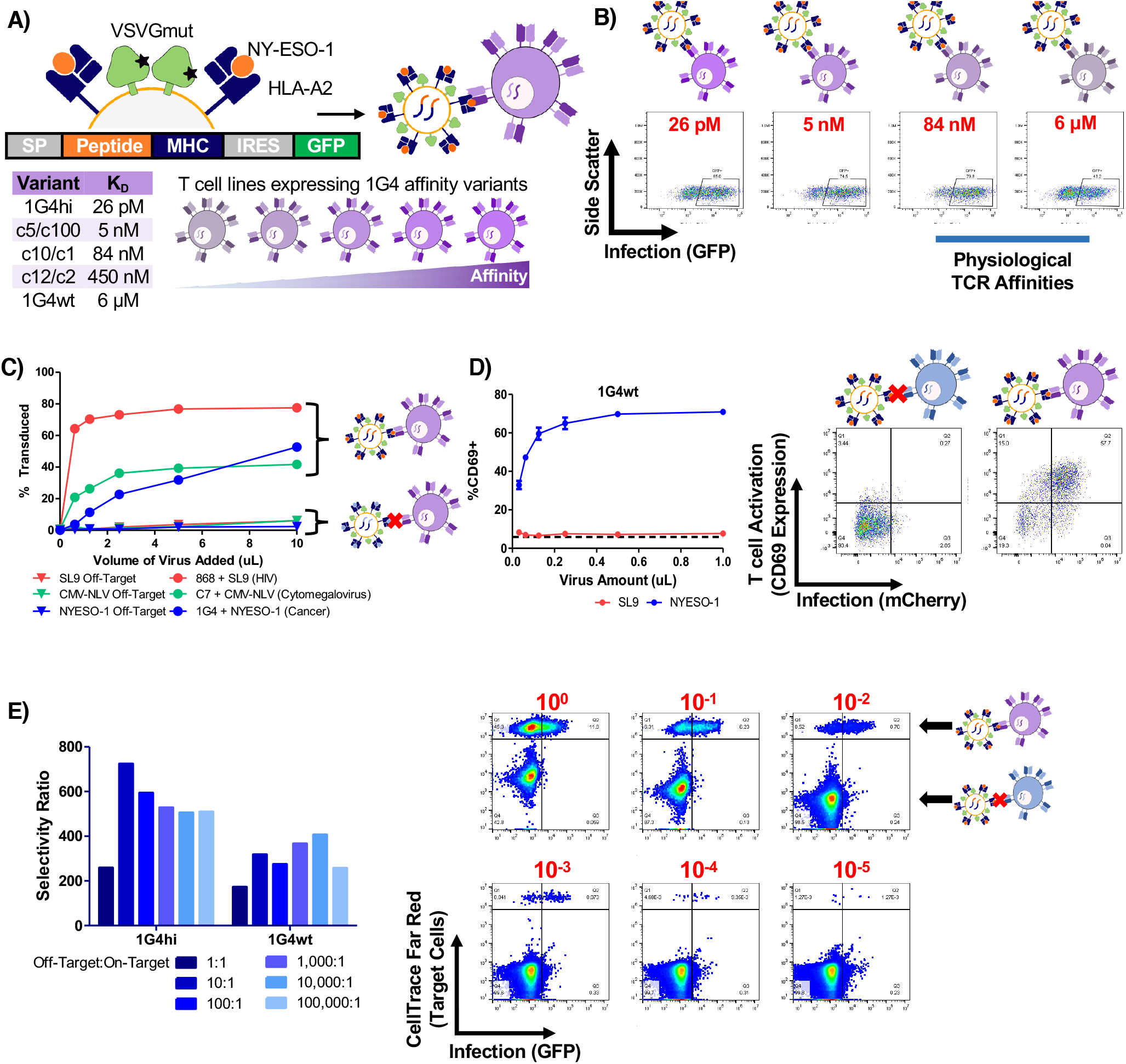
TCR-mediated infection is sensitive, specific, and induces signaling. **A**, Schematic of the 1G4 TCR-HLA-A2/NY-ESO-1 system used for TCR-pMHC characterization. **B**, Representative infection data for various 1G4 TCR affinity variants by HLA-A2/NY-ESO-1 displaying VSVGmut lentiviruses. **C**, infection of J76 cells via specific TCR-pMHC interactions via pMHC-displaying VSVGmut lentiviruses for three independent TCR-pMHC pairs (HLA-A2/SL9-displaying viruses infecting 868 TCR-expressing J76 cells; HLA-A2/NLV-displaying viruses infecting C7 TCR-expressing J76 cells; HLA-A2-NY-ESO-1-displaying viruses infecting 1G4 TCR-expressing J76 cells). **D**, Upregulation of CD69 on J76 cells transduced with the 1G4wt TCR during viral entry demonstrates that viral entry induces TCR signaling; data shown represent mean + SD across three biological replicates, dashed line represents CD69 expression in untreated J76-1G4wt cells. **E**, Selectivity ratio of on target to off target infection of J76 cells expressing 1G4 TCR variants mixed at indicated ratios with off-target Jurkat cells; representative of three independent experiments, flow plots correspond to 1G4wt TCR-expressing J76 cells with target cell frequencies indicated in red.

We next sought to determine the limitations of our system by characterizing its sensitivity and specificity using the 1G4-NY-ESO-1 system. To determine the ability of our approach to discern on-target interactions in complex mixtures, we labeled 1G4-expressing cells with a cell tracking dye, mixed them at varying ratios with unlabeled Jurkats, and infected the mixture with NY-ESO-1/HLA-A2 displaying viruses (**Figure 2E**). We calculated a selectivity ratio of on-target infection relative to off-target infection, and observed on-target selectivity greater than 200:1, down to target cell frequencies of 1×10^−5^. Thus, we concluded that our system can detect on-target interactions in the micromolar affinity range, even in complex mixtures containing hundreds of thousands of non-target T cells.

### Development of viral packaging strategy that maintains genotype-phenotype linkage

To fully enable interaction screening at library scale, we next turned to the challenge of producing the lentiviral libraries. While lentiviral screens have been widely used for functional genomics, standard lentiviral packaging techniques pose inherent limitations that are particularly relevant to interaction screens. The key challenge stems from the fact that at least hundreds of plasmids containing library elements are mixed in each transfected packaging cell (**Figure 3A**). In functional genomics, this leads to well-described intermolecular recombination of library elements^48,49^, which is a significant source of noise in combinatorial experiments. To date, solutions to this problem have included plasmid dilution (accompanied by a 100x reduction in viral titer)^49^, or simply restricting libraries to sizes suitable for arrayed screens. This problem is compounded in our approach, as multiple plasmids entering the same packaging cell compromises the link between viral genotype and surface phenotype.

**Figure 3.**
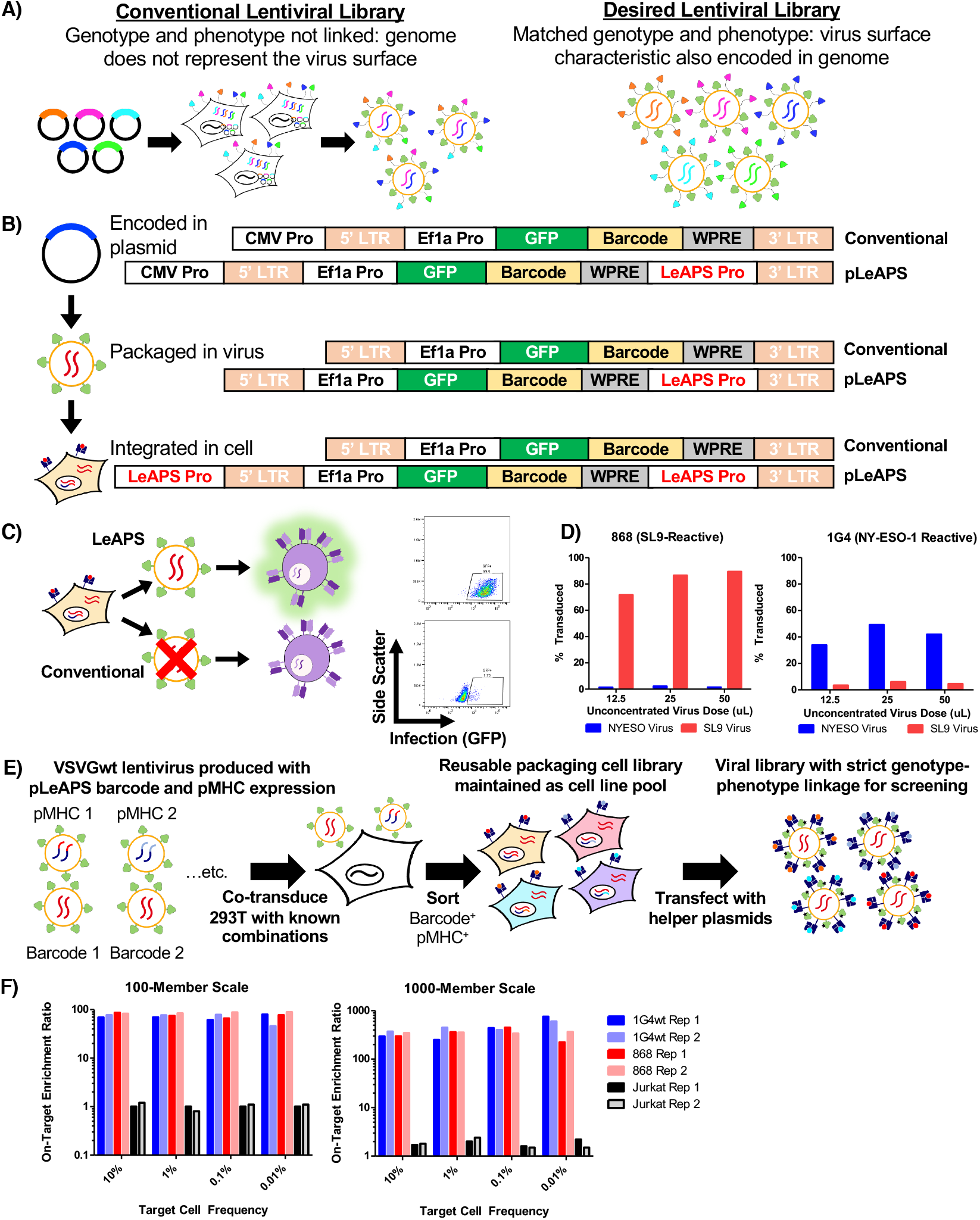
LeAPS-based lentiviral packaging enables scalable viral library construction. **A**, Schematic overview of the limitation of conventional lentiviral packaging strategies for surface displayed lentiviral libraries, in which a mixture of displayed surface proteins disrupts the linkage between viral pseudotype and genomic identity. **B**, Schematic overview of the mechanism of promoter translocation used in the LeAPS system, in which a promoter sequence placed internally in the lentiviral genome is placed 5’ of the 5’ LTR during the viral lifecycle, thus allowing the integrated vector to produce viral genomes at high copy number. If cells are infected with a single lentiviral vector, each infected cell is ensured to produce viruses that match viral surface phenotype and genotype. **C**, Packaging cells transduced with viruses containing LeAPS-encoding genomes and then transfected with lentiviral helper plasmids produce high titer virus (top), while those transduced with viruses containing standard genomes produce minimal functional virus (bottom). **D**, LeAPS-produced viruses maintain TCR specificity in two independent TCR-pMHC systems, demonstrating functional targeted viruses can be created. **E**, Schematic overview of the implementation of LeAPS for viral library assembly. 293T cells are co-transduced with known combinations of LeAPS-fluorophore-barcode viruses and pMHC expression viruses, all cells are pooled and sorted to generate a library packaging line that generates virus following helper plasmid transfection. **F**, On-target enrichment of libraries using defined mixtures of A2-NY-ESO-1 and A2-SL9 packaging cells to represent 100- or 1,000-variant libraries.

To obviate this issue, we exploited a detail of HIV-1 replication that results in the copying of sequences between the polypurine tract and the 3’ LTR to the 5’ end of the genome during reverse transcription and integration^50^. There have been several recent reports exploiting this phenomenon to enable genomic screens via copying of CRISPR gRNA cassettes^51,52^. Here, we insert a promoter to enable re-packaging of the viral genome following infection of a cell^53^ (**Figure 3B**). Conventional 3^rd^ generation lentiviral packaging approaches are inherently self-inactivating in part due to an inability to transcribe packageable viral genomes following infection; i.e. the LTRs, which are weak promoters, are truncated, and there is no promoter upstream of the viral genome following successful integration. Our approach, which we term **Le**ntiviruses **A**ctivated by **P**romoter **S**huffling (LeAPS), enables generation of lentiviruses from previously transduced cells upon the re-introduction of helper plasmids. As a result, we can use lentiviral transduction to ensure a single library member per packaging cell without incorporating additional viral elements into the genome. We first validated that viruses produced with the LeAPS strategy yielded high-titer virus (>10^6^ TU/mL unconcentrated), whereas conventional constructs (without a LeAPS promoter) yielded very little following transduction (**Figure 3C**). By stably expressing pMHC cassettes in LeAPS-transduced packaging cells, we verified that viruses produced in this manner maintain antigen-specific targeting capability (**Figure 3D**).

To develop a scalable library assembly strategy, we co-transduced 293T cells with two VSVGwt-pseudotyped viruses: one that drives surface expression of a known pMHC using a conventional lentivirus, and another that uses the LeAPS system to deliver a repackageable genome driving expression of a fluorescent protein and a defined barcode (**Figure 3E**). Following initial transduction, packaging cells for each library member were pooled and sorted for expression of pMHC and barcode in a single sorting step. This yields a library packaging line that can be used to generate the same lentiviral library indefinitely via single transfections with only helper plasmids. By contrast, conventional approaches require arrayed transfection to produce each library member for each experiment^54^, which poses a substantial impediment to experimental throughput.

To validate the utility of our approach, we employed a two-component system of the HIV SL9 antigen (SLYNTVATL) paired with a LeAPS-mCherry cassette and the NY-ESO-1 antigen paired with a LeAPS GFP cassette. By mixing the packaging cells for each at different ratios, we assessed the feasibility of pMHC libraries at 100-member and 1,000-member scales (**Supplementary Figure 3**). We then used flow cytometry to quantify signal to noise across different frequencies of on-target cells (**Figure 3F**). Across both library sizes, and extending from target cell frequencies of 10% down to 0.01%, we observed robust enrichment of on-target interactions, but minimal enrichment of off-target interactions (i.e. no enrichment of either component on Jurkat cells expressing an irrelevant TCR).

### Receptor-Antigen Pairing by Targeted Retroviruses

With all the necessary tools for RAPTR validated, we assembled a library of 96 pMHC constructs consisting of known seroprevalent viral or vaccine antigens from the IEDB^55^, and further filtered for binding to HLA-A2 using NetMHC4.1^56^ (**Table 1**). We co-transduced each into 293T cells alongside a LeAPS vector encoding a barcode, pooled the resulting cells proportionally, and sorted for cells expressing both barcode (via mCherry) and pMHC (via GFP). We verified that antigen and barcode expression in our packaging cell library was stable after at least 9 passages post-sort (**Supplementary Figure 4A**). We next used the library to infect a cell line expressing the C7 TCR^57^, which is known to recognize the CMV peptide NLVPMVATV (known as NLV). Without any cell sorting step required, we sequenced genomic DNA from transduced cells to enumerate barcode frequencies (**Figure 4A**). We observed a large enrichment of NLV barcode in transduced cells relative to the packaging cell line (**Figure 4B**), with NLV representing 15% of all reads in transduced cells and no notable enrichment of other sequences. We further validated this approach in C7 replicates (**Figure 4C** and **Supplementary Figure 4B**), and in a separate screen of the JM22 TCR ^57^, which recognizes the GL9 peptide from influenza A (**Figure 4D** and **Supplementary Figure 4C**). In all cases, we observed robust enrichment of on-target, but not off-target, barcodes.

**Table 1:** Summary of HLA-A2-restricted viral antigens included in RAPTR T cell library.

| Antigen Source | # of Epitopes |
| --- | --- |
| Adenovirus 5 | 3 |
| Herpes simplex virus 1 (HSV-1) | 31 |
| Herpes simplex virus 2 (HSV-2) | 4 |
| Varicella Zoster Virus (VZV) | 4 |
| Cytomegalovirus (CMV) | 6 |
| Coronavirus 229E | 2 |
| Coronavirus OC43 | 1 |
| Epstein-Barr Virus (EBV) | 26 |
| Influenza A (IAV) | 6 |
| Measles | 4 |
| Mumps | 6 |
| Kaposi's sarcoma-associated virus (HHV-8) | 3 |

**Figure 4.**
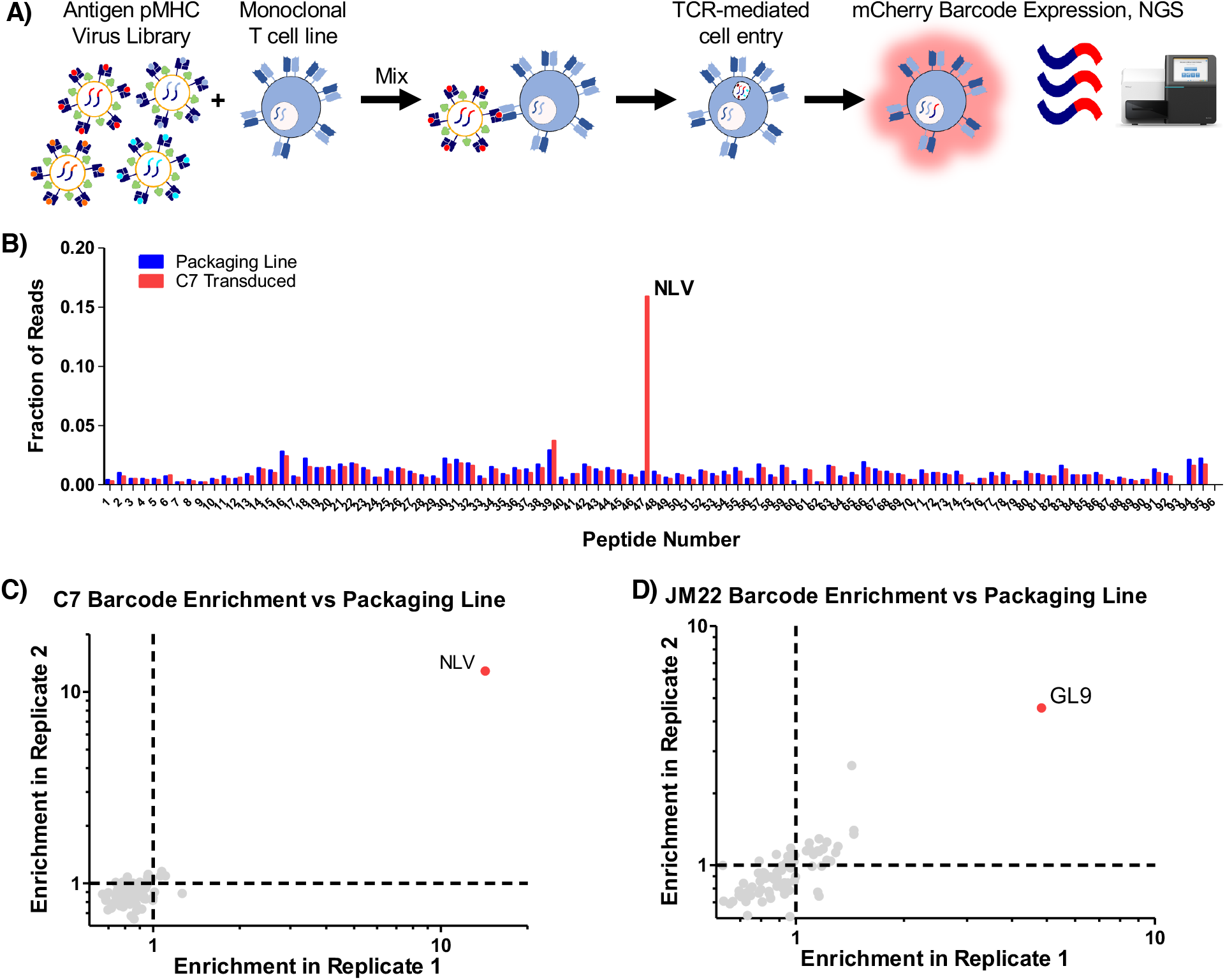
RAPTR enables facile antigen identification for T cell receptors. **A**, Workflow for RAPTR on monoclonal cell lines. **B**, Read fraction of barcodes following infection of C7 TCR-transduced J76 cells. **C**, Comparison of antigen enrichment upon library infection of C7 TCR-transduced J76 cells across two additional replicates. **D**, Comparison of antigen enrichment upon library infection of JM22 TCR-transduced J76 cells across two replicates.

### RAPTR can be adapted to BCRs

Many of the same constraints that limit TCR-pMHC interaction screens are also shared by efforts to identify B cell antigens at scale. Most approaches rely on barcoded multimers for very few targets, and the largest antigen library reported to date is 9 antigens^15^. As BCRs are also rapidly endocytosed upon antigen engagement^58,59^, we reasoned that we could use stabilized viral surface antigens or known immunogens as targeting ligands for viral infection. We thus generated a “hybrid” pseudotype co-expressing either the surface-tethered receptor binding domain (RBD) or the full-length, pre-fusion stabilized spike (S2P) protein from SARS-CoV-2^60^ alongside VSVGmut (**Figure 5A**). We used each of these to achieve efficient, antigen-specific infection of a Ramos cell line expressing CR3022, a B cell receptor clone that is cross-reactive for the RBDs of SARS-CoV-1 and 2^61,62^ (**Figure 5B**). Similarly, hybrid pseudotypes incorporating HIV env CD4 binding site (CD4bs) constructs from diverse clades^63^ each infected cell lines expressing the bNab VRC01^64^, and a known affinity-reducing mutation (D368R) dramatically reduced, but did not fully abrogate, infection^65^.

**Figure 5.**
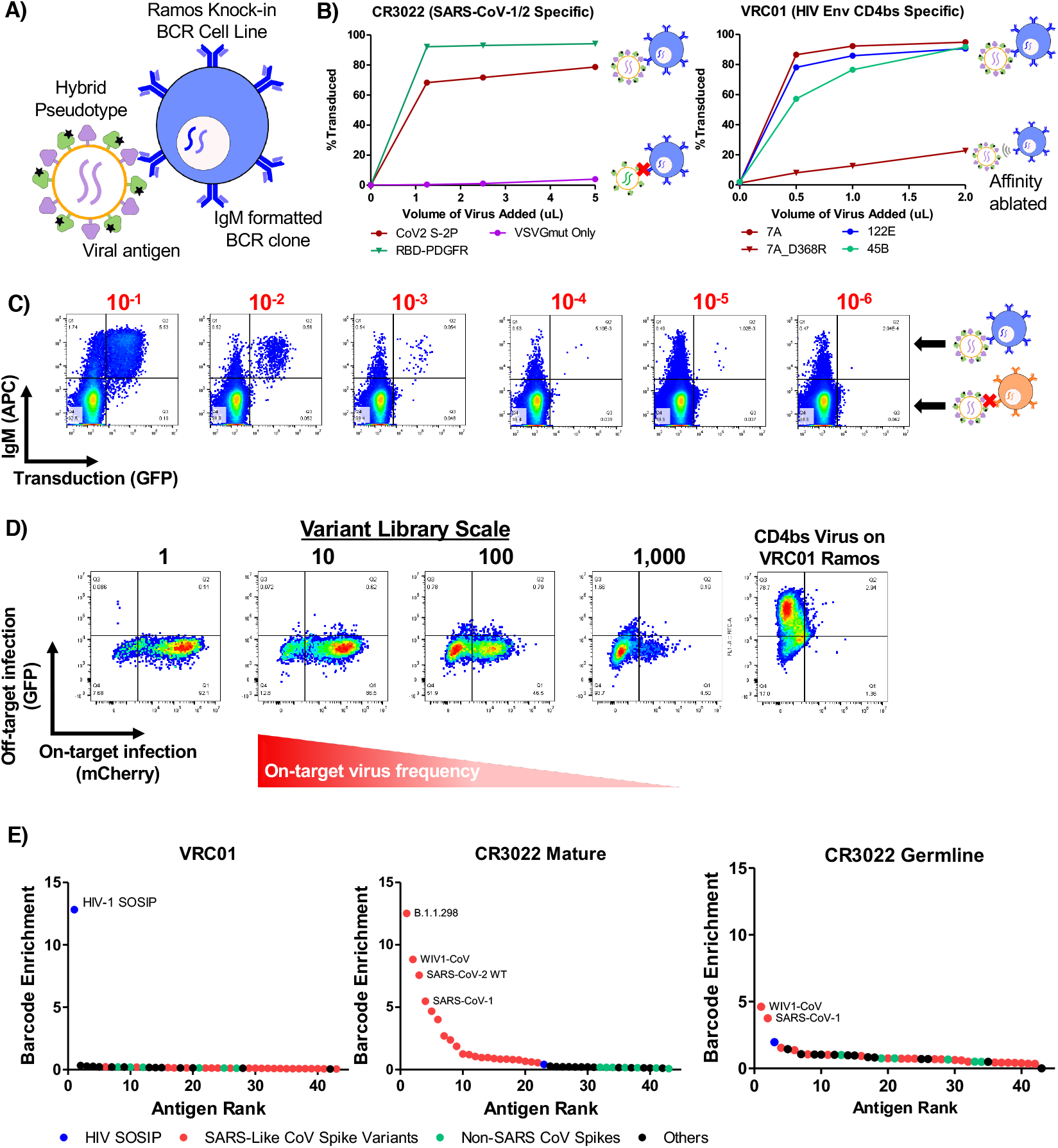
RAPTR enables profiling of B cell reactivity via infection of BCR knock-in cells. **A**, Overview of BCR-based targeting via the VSVGmut hybrid pseudotype in which a viral protein is co-expressed with VSVGmut. **B**, Targeting of Ramos cells expressing the SARS-CoV S protein specific CR3022 BCR via VSVGmut lentiviruses displaying either SARS-CoV-2 spike (2P) or surface-bound SARS-CoV-2 S protein RBD, or VRC01 BCR via env constructs representing multiple clades of HIV; representative data from 2 independent experiments. **C**, Infection of CR3022 BCR-expressing Ramos cells mixed into IgM-negative Ramos cells at target cell frequencies indicated in red, demonstrating selectivity in a mixed population of cells. **D**, Infection of CR3022 BCR-expressing Ramos cells with mixtures of SARS-CoV-2 RBD (on-target, expressing mCherry) and HIV env (off-target, expressing GFP) viruses; variant library size indicates the ratio of off-target to on-target virus present. Infection of VRC01-expressing Ramos cells with HIV env hybrid pseudotyped viruses (right) demonstrates that the viral particles remain functional. **E**, Enrichment of antigen barcodes following viral glycoprotein library infection of Ramos cells expressing BCRs for VRC01, mature CR3022, or the germline-reverted version CR3022 demonstrating selectivity even for antibody sequences before affinity maturation.

To determine the sensitivity for BCR-mediated cell entry, we mixed Ramos cells expressing either the CR3022 BCR (IgM^+^) or no BCR (IgM-negative), and tracked on-target cells via surface IgM expression (**Figure 5C**). Using RBD-displaying viruses, we observed specific infection at target cell frequencies at least as rare as 1×10^−5^, but could not calculate a meaningful selectivity ratio due to a lack of off-target infection. Next, to assess the potential feasible library size, we mixed on-target RBD-displaying viruses encoding GFP with off-target CD4bs-displaying viruses encoding mCherry. and used them to infect CR3022 BCR-expressing Ramos cells (**Figure 5D**). Keeping the total virus amount constant, we were able to robustly detect on-target interactions with minimal off-target infection in mixtures as dilute as 1:1,000 on-target viruses. We observed similar results when displaying SARS-CoV-2 S2P instead of RBD (**Supplementary Figure 5**). These results gave us confidence in the feasibility of extracting meaningful signal from interaction screens using lentiviral infection via the BCR.

Next, we sought to establish the capabilities of RAPTR for BCR profiling in a true library setting. We assembled a library composed of 21 prefusion-stabilized SARS-CoV-2 spike variants and 22 additional vial surface glycoproteins or immunogens from diverse sources as controls to enable proof of concept experiments (**Table 2, Supplementary Table 2**). When we used this library to infect cells expressing the VRC01 BCR, we observed clear enrichment of only HIV-1 SOSIP^66^ (**Figure 5E**), highlighting the potential for direct receptor deorphanization. When we infected cells expressing the CR3022 BCR, we observed clear enrichment of SARS-CoV spikes, including multiple SARS-CoV-2 variants. Notably, a version of CR3022 that excluded the mutations acquired via somatic hypermutation yielded enrichment of only SARS-CoV-1 spike and the closely related WIV-1 spike, highlighting the ability of RAPTR to distinguish BCR cross-reactivity even for naïve BCRs that have yet to undergo affinity maturation.

**Table 2:** Summary of viral surface glycoproteins for RAPTR B cell library.

| Antigen Source | # of Variants |
| --- | --- |
| SARS-CoV-2 Spike | 21 |
| Other MERS- and SARS-like CoV Spike | 4 |
| Common Cold CoV Spike | 4 |
| SARS-CoV-2 E/M | 2 |
| Influenza | 4 |
| Seroprevalent or Childhood Vaccine Viruses (Positive Controls)<br>Measles, RSV, CMV, EBV | 5 |
| Emerging or Uncommon in Healthy Donors (Negative Controls)<br>HIV, Dengue, Nipah Virus | 3 |

### RAPTR scales to library-on-library screens

To fully realize the potential of RAPTR for antigen identification, we sought to apply it to library vs library screening by using our 96-member pMHC viral library to pan a previously reported library of >450,000 TCRs^67^. To simplify the first attempt, we pre-enriched for potentially reactive cells using tetramers for CMV, EBV, and Flu (CEF) antigens presented by HLA-A2, resulting in a polyclonal pool of TCRs with greater prevalence of these specificities, similar to the routine process of pre-expanding cells (**Figure 6A**). Following transduction and FACS to isolate transduced cells, we used bulk sequencing to identify which antigens were enriched in aggregate. In line with tetramer staining of the untransduced cells, we observed strong enrichment of Flu GL9, and more modest enrichment of EBV GLC (**Figure 6B, Supplementary Figure 6**).

**Figure 6.**
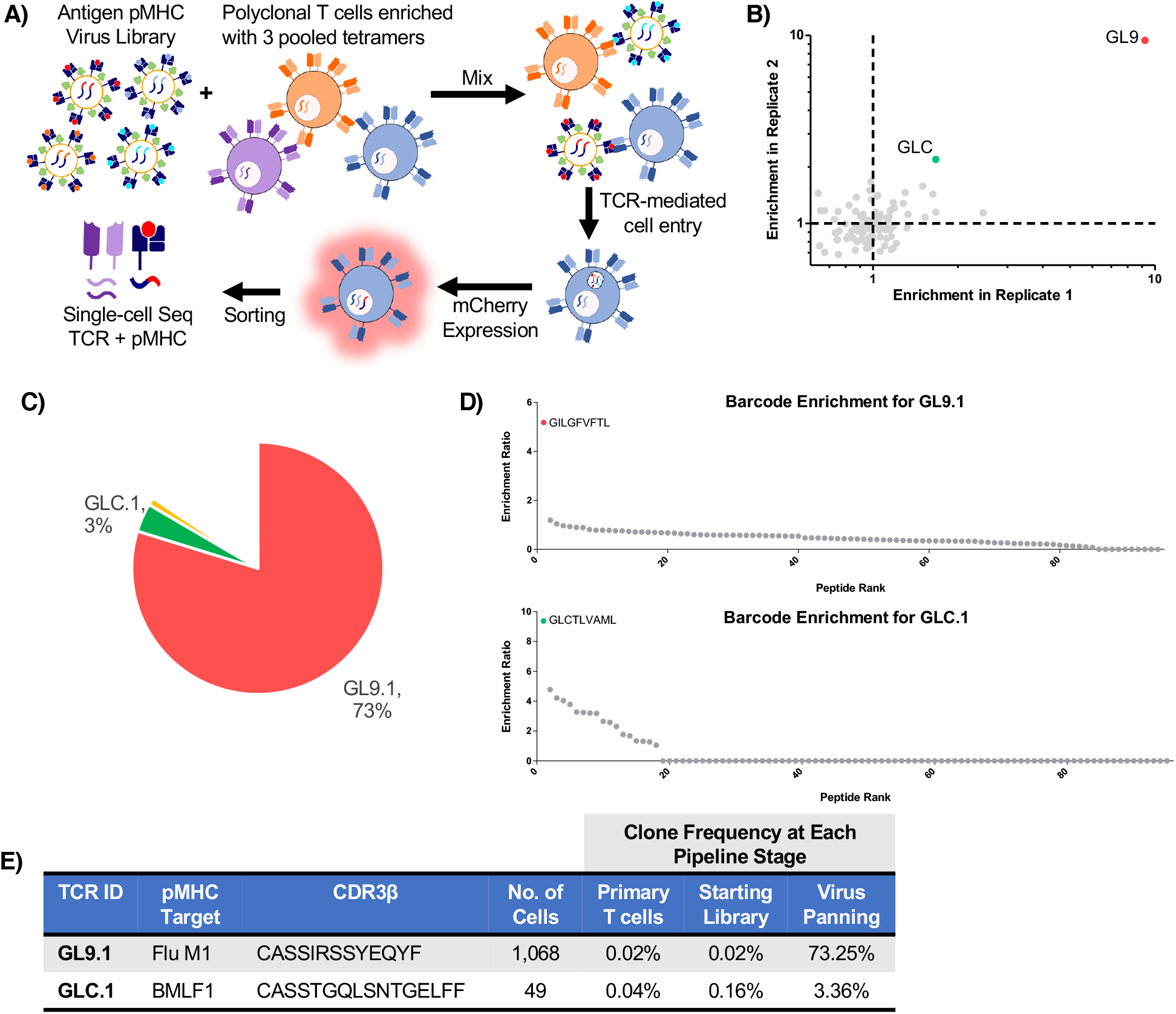
RAPTR scales to polyclonal antigen identification in a single, scalable pipeline. **A**, Schematic overview of the library on library RAPTR paradigm for TCR antigen identification. **B**, Bulk sequencing of barcodes following infection of enriched CSS-930 TCR library with viral antigen library shows enrichment of immunodominant EBV and Influenza antigens in two independent infections. **C**, Clonal frequencies of enriched TCRs in cells analyzed by single cell sequencing; white space represents TCRs found in < 2 cells. **D**, Enrichment of antigen barcodes in the top two most abundant clones. **E**, Detailed tracking of TCR clones with matched antigens, including CDR3 sequence, number of cells analyzed, and clonal frequencies at each stage of the pipeline.

With this validation in hand, we performed single-cell RNA sequencing using the 10x Genomics Chromium 5’ chemistry with V(D)J enrichment. We included a targeted primer at the reverse transcription step and a custom amplification protocol to enable recovery of our pMHC barcodes, akin to ECCITEseq and other similar methods^68,69^. In this experiment, we recovered 1,458 cells, with TCR clonotypes dominated by a few clones (**Figures 6C** and **6E**). We grouped cells of common clonotype together and analyzed pMHC barcode expression to determine enrichment relative to packaging library frequency. The most prevalent TCR clone, GL9.1, demonstrated strong enrichment of GL9 pMHC signal (**Figure 6D**). Upon literature search, we noted that GL9.1 represents a well-characterized public clonotype known to recognize GL9 peptide presented by HLA-A2^70^.

The next most abundant clone, GLC.1, was identified in a previous screen of this same TCR library by recombinant HLA-A2 GLC dextramer^67^. Our screen also identified GLC as the target pMHC (**Figure 6D**). Notably, this was the only clone previously identified by GLC dextramer panning that was validated by monoclonal binding and functional tests. Our approach did not identify any of the previously reported false positives as hits, providing further evidence that RAPTR is an efficient means of identifying T cell antigens that integrates both binding and signaling.

## Discussion

RAPTR is a high-throughput platform for directly linking immune receptors with their cognate antigens. It is based on the combination of several conceptual and technological advances that enable (i) versatile and efficient targeted viral entry via the VSVGmut pseudotyping system; (ii) a method for scalable, reproducible lentiviral library packaging (LeAPS); and (iii) the use of viral entry as a means of screening for interactions. We demonstrated a system for efficient, antigen-specific infection via both the TCR and the BCR that integrates both binding and target cell activation. We then exploited this mode of infection to match receptors with target antigens in complex mixtures and in a library vs library format.

We demonstrated that the VSVGmut pseudotyping system is efficient, specific, and uniquely modular by targeting a variety of cell surface proteins, including cytokine receptors, costimulatory molecules, lineage markers, and both T and B cell receptors. We report here as many different targeting approaches for VSVGmut as have been reported for any other single pseudotyping system^27^, including the first true antigen-specific infection of lymphocytes and the first combinatorial targeting approach. In addition, the VSVGmut system produces viruses at higher average titers than other systems while maintaining a high degree of specificity. Whereas most pseudotyping systems have been limited to the use of small, stable, high-affinity targeting ligands (e.g. scFvs or DARPins), VSVGmut is uniquely versatile in ligand usage due to the modular design it permits. We report targeting even at micromolar affinities, whereas the efficiency of infection by paramyxoviral pseudotypes was reported to dramatically decrease in the low nanomolar range^71^.

### Applications of RAPTR

We have demonstrated the utility of RAPTR for identifying antigens recognized by TCRs and BCRs. For T cell reactivity, our approach could be directly applied to rapidly deorphanize TCRs from existing single-cell studies, which routinely pair TCR sequences with gene expression for cancer, autoimmunity, or infectious disease. Existing techniques, including yeast display, SABRs, and T-Scan, require individual screens to pair each TCR with its cognate antigen, including at least one cell sorting step per TCR. In contrast, RAPTR can be performed on pools of TCRs in a single step, with only one cell sorting step required (and no sorting required for single-TCR experiments). While barcoded pMHC multimers can be built in to single-cell workflows, large antigen pools are inherently difficult to produce and validate, and require equivalent effort for each new batch. These factors have limited their use to only a few specialized labs thus far. RAPTR library production is straightforward and far more scalable in comparison, requiring only a simple transfection per batch following initial library assembly. In principle, these reagents could be made and distributed at large scale, either via sharing the packaging cell line or by leveraging current industrial infrastructure for production of lentiviruses.

For BCRs, we envision applying pools of protein variants, such as the one presented here, to large libraries of BCRs. Such an approach would greatly accelerate isolation and optimization of cross-reactive mAbs to a wide range of target classes. Current techniques require the recombinant expression of individual mAbs followed by ELISA for each target of interest, which represents a major bottleneck to experimental scale and scope^72^. Using RAPTR with BCR knock-in cells, which could themselves be prepared as pooled lentiviral libraries^62^, could alleviate this bottleneck. For example, even scales smaller than the one we presented would be sufficient to represent every subtype of influenza hemagglutinin and neuraminidase, multiple representatives of each of the major clades of HIV-1, and more. Larger libraries on the scale of 1000s of variants, whose feasibility we demonstrated, could enable detailed antigenic site or epitope mapping for many mAbs at once, a process currently limited to structural studies that are difficult to scale.

Finally, the demonstrated modularity of the system opens the door to many potential applications. Since our approach only requires 293T cell surface expression of the targeting molecule, we hypothesize that RAPTR can be readily adapted to other class I MHC alleles, class II MHC alleles, as well as non-human systems for vaccine and immunotherapy development. This compares favorably to recombinant MHC expression or yeast display, each of which can require allele-specific optimizations and/or mutations to ensure proper folding^8,73,74^. However, our system is not restricted to immunological applications. In principle, it is applicable to any receptor-ligand system in which one binding partner can be displayed on a lentivirus and the other can be expressed as an endocytic receptor on cells. This may enable large-scale interactome and co-evolution studies that are not readily achievable by existing techniques. Furthermore, the VSVGmut pseudotyping system may be useful as a new platform for the development of efficient, specific gene therapies.

### Limitations

As with any technology, RAPTR has several limitations. Our proof of concept studies were performed on ∼100-antigen scale, and while our control experiments indicate this could be readily increased to 1,000s, these scales will still require pre-selection of antigen targets for analysis. Future work will focus on further streamlining library assembly and increasing transduction efficiency toward larger-scale assays. In addition, the single-cell sequencing step must be performed in sufficient depth to facilitate acceptable signal to noise determinations. While we chose to use a commercially available platform to ensure broad applicability, recent and future advances in the scale of single-cell analysis^75^ will improve the utility of RAPTR regardless of platform. Like other genetically encoded techniques, RAPTR also requires prior selection of MHC haplotype(s) to generate libraries. Finally, the antigen identification studies we presented were on cell lines with receptors knocked in, rather than primary cells. While the decreasing cost and turnaround time of gene synthesis will make such resources increasingly available, future studies can apply RAPTR directly to primary cells.

Altogether, RAPTR is a versatile platform for high-throughput interaction screens that uniquely incorporates diversity of both receptors and ligands in a single assay. The resulting tools will enable detailed studies of antigen recognition for understanding and engineering the adaptive immune response, as well as broader interactome studies.

## Methods

### Media and Cells

HEK293T cells (ATCC CRL-11268) were cultured in DMEM (ATCC) supplemented with 10% fetal bovine serum (FBS, Atlanta Biologics) and penicillin-streptomycin (Pen/Strep, Gibco).

Jurkat (ATCC TIB-152), J76 cells, and Ramos cells (ATCC CRL-1596) were cultured in RPMI-1640 (ATCC) supplemented with 10% FBS and Pen/Strep. J76 cells^76^ (PMID 29707134) were a gift from Mirjam Heemskerk and Mark Davis.

CSS-930 TCR library cells were a gift from David Johnson^67^, and were cultured in RPMI-1640 (ThermoFisher) supplemented with 10% FBS, Pen/Strep, 1x non-essential amino acids (ThermoFisher), 2mM GlutaMAX (ThermoFisher), and 1mM sodium pyruvate (ThermoFisher).

### Plasmid Construction

The plasmid pHIV-EGFP was gifted by Bryan Welm & Zena Werb (Addgene plasmid #21373) and pMD2.G and psPAX2 were gifted by Didier Trono (Addgene plasmid #12259 and #12260). pLentiCRISPR v2 was a gift from Feng Zhang (Addgene plasmid # 52961). IL-13 (Uniprot ID P35225) residues 35-146 were cloned into the pHIV backbone containing the Igκ leader peptide, the PDGFR stalk and transmembrane domain (Uniprot P09619 residues 449-497) with extracellular linkers listed in Supplementary Figure 1B. The anti-CD19 scFv FMC63 and was cloned into the IgG4 hinge-PDGFR display format in the pMD2 backbone. The anti-CD3 Fab UCHT1 was cloned into the PDGFR stalk only format. Human CD80 (Uniprot P33681, residues 1-273) was cloned into the pMD2 backbone. pMHC single-chain trimers^42^ were cloned into either the pMD2 backbone for individual infections or the pHIV backbone for library construction. To generate the pLeAPS backbone, the CMV core promoter was cloned into the pLenti backbone between the polypurine tract and the 3’ LTR, analogous to previous work^52^. For individual infections, SARS-CoV-2 RBD or HIV env CD4bs constructs were cloned into the pMD2 backbone on the PDGFR stalk only display architecture. Prefusion-stabilized SARS-CoV-2 spike (2P) was cloned into the pMD2 backbone for individual infections. For library assembly, viral constructs described in **Supplementary Table 2** were cloned into the pLenti backbone, C-terminally fused to GFP via a P2A motif. For pLeAPS barcodes, mCherry with an 8-nt degenerate sequence was cloned into the pLeAPS backbone downstream of the Ef1α core promoter.

### Transfection for Lentiviral Production

Lentiviruses were prepared by transient transfection of HEK293T cells with linear 25 kDa polyethylenimine (PEI, Santa Cruz Biotechnology) at a 3:1 mass ratio of PEI to DNA. Briefly, DNA and PEI were diluted in Opti-MEM (ThermoFisher) and mixed to form complexes. Complex formation was allowed to proceed for 15 minutes at room temperature before dropwise addition to cells. Media was changed to complete DMEM + 25 mM HEPES after 3-6 hours.

For individually targeted or VSVGwt viruses, plasmid mass ratios were 5.6:3:3:1 for transfer plasmid to psPAX2.1 to targeting plasmid (when used) to fusogen plasmid (either VSVGmut or VSVGwt). Targeting plasmids contain expression cassettes for virally-displayed ligands in the pMD2 backbone. Total plasmid amounts are indicated in the table below. For LeAPS-based virus production, packaging cells were transfected with a 3:1 mass ratio of psPAX2.1:fusogen.

| Production Scale (per well) | Total Plasmid Amount (µg) |  |  |
| --- | --- | --- | --- |
|  | Non-Targeted | Targeted | LeAPS-based |
| 6-well plate | 2.4 | 3.15 | 1 |
| 10 cm dish | 24 | 31.5 | 10 |
| T225 | 72 | 94.5 | 30 |

### Viral Purification

Unconcentrated viruses were filtered (0.45μm PES) and used directly. If needed, they were stored at 4°C for up to 2 weeks or at -80°C indefinitely. Concentrated viruses were filtered (0.45μm PES) and concentrated 200x by ultracentrifugation for 90 minutes at 100,000xg at 4°C. The supernatant was discarded and viral pellets were resuspended in Opti-MEM overnight at 4°C.

### Single lentiviral infections

Individual infections were carried out with the indicated amounts of virus and cells, in the presence of 8 μg/mL of either polybrene or diethylaminoethyl (DEAE)-dextran (Sigma-Aldrich). After 24h, an additional 1x volume of media was added. Cells were analyzed by flow cytometry 48-72h post-infection for Jurkat or J76 infections, and 24-48h post-infection for Ramos cell infections. If cells were only assessed for viral infection, cells were washed once in FACS buffer (PBS + 0.1% BSA and 1 mM EDTA) before analysis on an Accuri C6 or Cytoflex S flow cytometer. In experiments where an additional cell lineage or activation marker such as CD25 or CD69 were assessed, cells were washed once in FACS buffer, stained for 10 minutes in FACS buffer containing a marker-specific antibody, then washed 2x with FACS buffer before analysis for infection (GFP or mCherry) via flow cytometry.

### Human Primary T cell activation and transduction

Peripheral blood mononuclear cells from healthy donors were purified from leukopaks purchased from Stem Cell Technologies using Ficoll-Paque PLUS (GE Healthcare) density gradient centrifugation with SepMate tubes (Stem Cell Technologies) per manufacturer instructions. Primary CD8^+^ T cells were isolated using EasySep Human CD8^+^ T Cell Enrichment Kits (Stem Cell Technologies) and cultured in RPMI-1640 (ATCC) supplemented with 10% fetal bovine serum, 100 U/ml penicillin-streptomycin (Corning), and 30 IU/ml recombinant human IL-2 (R&D Systems). Prior to transduction with VSVGwt viruses, T cells were activated using a 1:1 ratio of DynaBeads Human T-Activator CD3/CD28 (Thermo Fisher) for 24 hours, after which 8 µg/mL of polybrene (Santa Cruz Biotechnology) and concentrated lentivirus were added to culture at a multiplicity of infection of 1. For targeted viruses, the same protocol was used, but DynaBeads were omitted.

### Antigen receptor cell line generation

Lentiviral TCR cassettes were formatted as TCRβ-P2A-TCRα and cloned into the pHIV backbone. TCR KO J76 cells were transduced as described above and sorted based on TCR expression to establish monoclonal cell lines. Ramos BCR cells were established according to the published protocol^62^.

### Lentiviral infections in Mixed Cell Populations

Cells were labeled with CellTrace dyes (ThermoFisher) according to the manufacturer’s instructions, counted, and mixed at the indicated ratios. After labeling, viral infections were carried out as indicated above. After 48 hours, cells were washed in FACS buffer and analyzed via flow cytometry to examine infection in CellTrace^+^ vs CellTrace^-^ cells. After flow cytometry, the selectivity was calculated as follows:

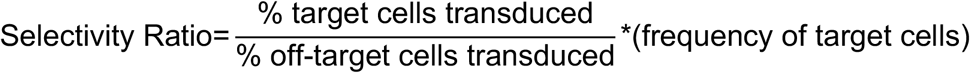

### Generation of LeAPS libraries

To generate LeAPS production cell lines, 293T cells were transduced as outlined above while being seeded at 20% confluency on either 6-well or 96-well plates (ThermoFisher). For 6-well plates, 500 μL each of unconcentrated, VSVGwt-pseudotyped LeAPS barcode and ligand expression viruses were used for infection. 50 μL of each virus was used for 96-well plates.

For libraries, cells were transduced in duplicate. One of the duplicates was assessed by flow cytometry to determine the proportion of cells transduced with both ligand (GFP^+^) and LeAPS barcode (mCherry^+^). Cells from the remaining duplicate were then pooled, with each library member normalized to include an equivalent number of ligand-expressing (GFP^+^) and LeAPS barcode-transduced (mCherry^+^) cells for each library member. This pool of cells was then sorted based on GFP and mCherry expression to make library packaging cell pools. To produce lentiviruses, library packaging cells were transfected as described above, but using only psPAX2.1 and pMD2-VSVGmut. For library creation, both transfer plasmids and the pMD2 targeting plasmid were omitted, as they were replaced by the LeAPS packaging line. After 48 hours, virus was then collected and concentrated as described above.

### Library screening of monoclonal TCR lines

10 μL of concentrated pMHC library virus was used to transduce one million TCR-expressing J76 cells as outlined above. After infection was confirmed by flow cytometry as described above, genomic DNA was isolated using the PureLink Genomic DNA kit (ThermoFisher). Barcode inserts were then amplified via 25 cycles of PCR and submitted for Amplicon-EZ analysis by Genewiz. Enrichment was calculated for each barcode as the fraction of total barcode-containing reads divided by the barcode frequency in the packaging cells.

### Library screening of monoclonal Ramos BCR lines

500 μL of unconcentrated viral antigen library was used to transduce one million BCR-expressing Ramos cells as outlined above. After infection was confirmed by flow cytometry, genomic DNA was isolated using the PureLink Genomic DNA kit. Barcode inserts were then amplified via 25 cycles of PCR and submitted for Amplicon-EZ analysis by Genewiz or analyzed via a MiSeq v3 150×150nt PE Nano kit (MIT BioMicroCenter). Enrichment was calculated for each barcode as the fraction of total barcode-containing reads divided by the barcode frequency in the packaging cells.

### Tetramer enrichment of TCR libraries

To enrich a polyclonal pool of cells from the CSS-930 TCR library for known specificities, 50 million library cells were co-stained with an anti-TRBC antibody (clone IP26) and a pool of HLA-A2 tetramers (NIH Tetramer Core) presenting the following peptides: NLVPMVATV (NLV), GILGFVFTL (GL9), and GLCTLVAML (GLC), all at 1:800 dilutions. Sorting for double-positive cells was performed on a Sony MA900 FACS.

### Library vs library TCR-pMHC screen

10 μL of concentrated pMHC library virus was used to transduce two million tetramer-enriched cells. Cells were sorted for infection based on mCherry expression, and then submitted for analysis using the 10x Genomics Chromium 5’ v2 V(D)J kit with a barcode construct-specific primer spiked in prior to droplet encapsulation (see Supplementary Methods for further details). TCR amplicons were prepared and sequenced according to the manufacturer’s instructions. Following cDNA amplification, mCherry barcodes were enriched via separate PCRs and sequenced on an Illumina MiSeq (150×150nt paired end reads). CellRanger was used to assign TCR clone identities for each cell. Cell barcodes were used to match TCRs with associated pMHC barcodes counts.

## Data Availability

The next-generation sequencing datasets generated during and/or analyzed during the current study are available in the NCBI sequence read archive (accession numbers to be assigned before publication). All data generated or analyzed during this study are included in this published article, the sequence read archive, and its supplementary information files.

## Code Availability

A sample R workspace for extracting barcode counts based on CITEseq^68^ will be posted to Birnbaum lab code repository upon publication.

## Acknowledgments

We would like to thank the Koch Institute Swanson Biotechnology Center for their technical support, especially the Flow Cytometry Facility, MIT BioMicro Center, and High Throughput Sciences Facility. We thank Charlie Whittaker of the Barbara K. Ostrom (1978) Bioinformatics and Computing Core Facility for helpful discussion and implementation of single-cell sequencing data analysis pipelines. We also thank Glenn Paradis and Stuart Levine for many helpful discussions and suggestions. We thank Aaron Winkler, Kellie Kravarik, and Marc Wadsworth for helpful discussions and suggestions. We thank the NIH Tetramer Core Facility (contract number 75N93020D00005) for providing pMHC tetramer reagents used in this study.

This work was funded by grants from the Koch Institute Frontier Award program, the Packard Foundation, the Damon Runyon Cancer Research Foundation, the Michelson Medical Research Foundation, Pfizer, Inc., and the Department of Defense (W81XWH-18-1-0208) to M.E.B.; the National Institutes of Health (DP2AI158126 to M.E.B. and R01AI137057, R01AI153098 to D.L.); National Science Foundation Graduate Research Fellowship and Siebel Scholarship to C.S.D.; a Canadian Institutes of Health Research Doctoral Foreign Study Award to S.G.; a graduate research fellowship from the Ludwig Center at MIT’s Koch Institute to E.J.K.; and a Medical Scientist Training Program grant (T32 GM007753) from the National Institute of General Medical Sciences at Harvard Medical School to B.E.S.. M.E.B and S.K.D. were funded by a Technology Impact Award from the Cancer Research Institute and are Pew-Stewart Scholars in Cancer Research. Core facilities in the Koch Institute are partially by Cancer Center Support (core) Grant P30-CA14051 from the NCI.

## Author contributions

Conception of project: C.S.D., M.E.B. Conducting experiments: C.S.D., A.R., S.G., B.E.S., E.J.K., J.D., L.R., V.O. Data analysis: C.S.D., A.R., S.G., B.E.S., E.J.K., J.D. Supervision: D.L., M.D., S.K.D., M.E.B. Writing manuscript: C.S.D., M.E.B. Editing manuscript: all authors.

