## Supplemental Figures 1-7, Supplemental Tables 1-2, and Supplemental Methods for "Antigen identification and high-throughput interaction mapping by reprogramming viral entry"

A)

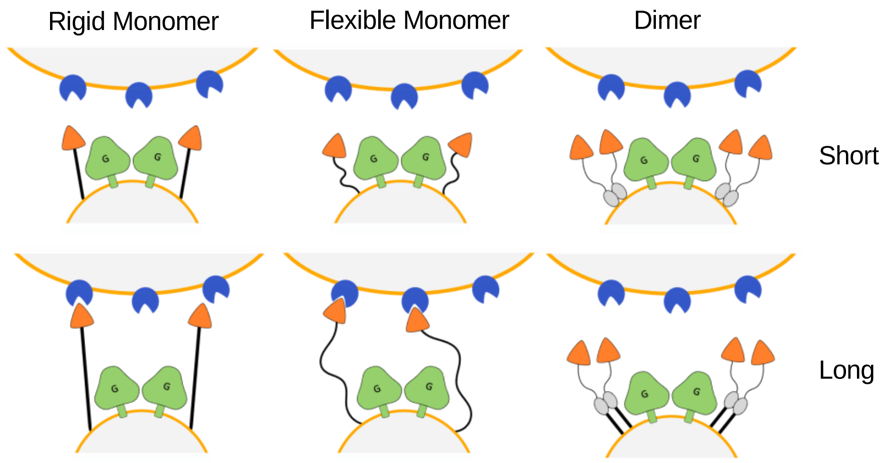


B)

| Architecture Type | Name | Unconcentrated Titer ($10^6$ TU/mL) |
|---|---|---|
| Short, rigid monomer | PDGFR transmembrane | 0.01 |
| Short, flexible monomer | GAPGAS linker | <0.01 |
| Short dimer | IgG4 hinge only | 0.01 |
| Extended, rigid monomer | PDGFR transmembrane and stalk | 0.046 |
| Extended, flexible monomer | $(G_4S)_3$ linker | <0.01 |
| Extended dimer | IgG4 hinge + PDGFR stalk | 0.83 |
| Extended dimer | CD8α stalk | 0.85 |
| Tetramer | Tetrameric coiled coil | 0.35 |

C)

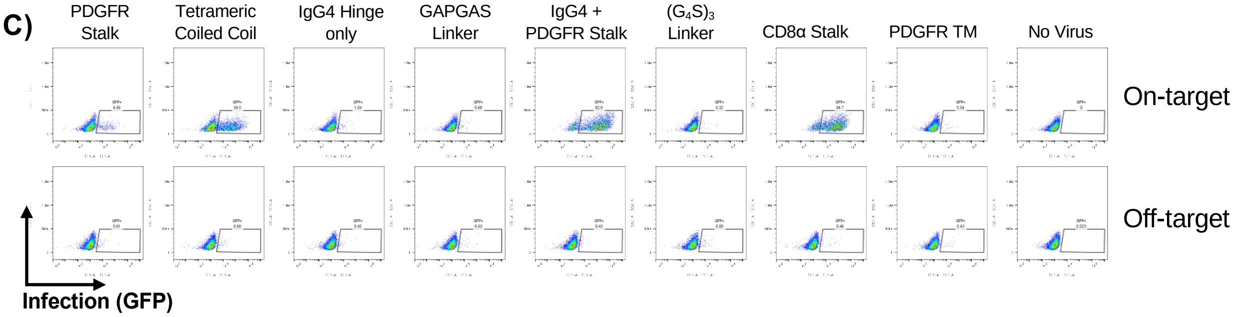


**Supplementary Figure 1 | IL-13 Surface Architecture Development**

**A,** Schematic diagram of the different targeting ligand architectures used. **B,** Summary table with details of each individual construct and titer achieved. **C,** Flow cytometry plots showing infection of IL13Rα1-expressing Jurkat cells (top, on-target) and parental Jurkat cells which don’t express the receptor (bottom, off-target)

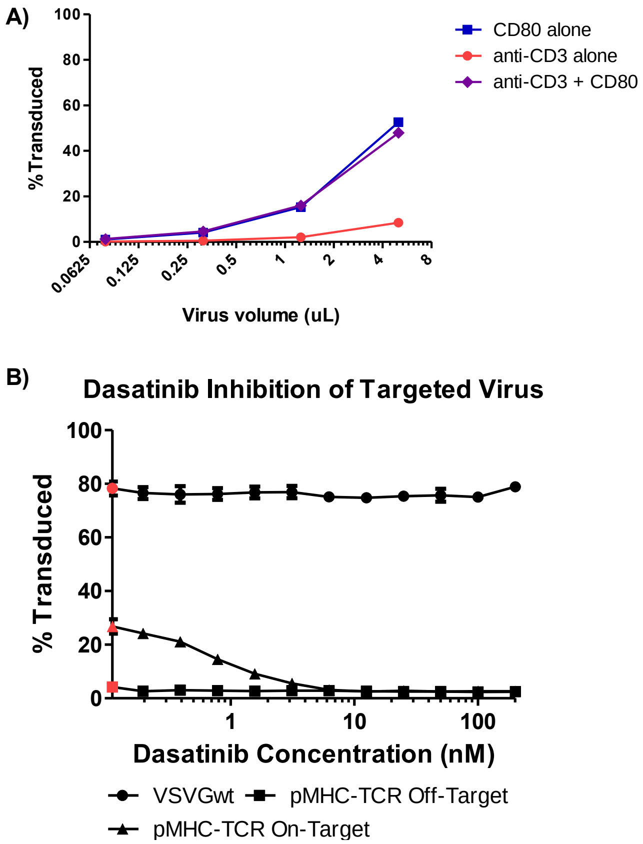


**Supplementary Figure 2 | Viral entry via the TCR complex and costimulatory receptors, and inhibition by dasatinib**

**A,** $1x10^5$ Jurkat cells were infected with the indicated amount of VSVGmut pseudotyped virus displaying either anti-CD3 Fab, CD80, or a combination of the two; data shown as mean + SD of three biological replicates **B,** $5x10^4$ J76 cells expressing the 868 TCR were pre-incubated with the indicated concentrations of dasatinib for 2 hours to inhibit TCR signaling prior to virus addition (red dots indicate no dasatinib condition). Next, 5 µL of virus displaying either VSVGmut and HLA-A2-SL9 (on-target), VSVGmut and HLA-A2-NY-ESO-1 (off-target), or VSVGwt were added. Infection was measured 48h post-infection. Results represent the mean and standard deviation of 3 biological replicates.

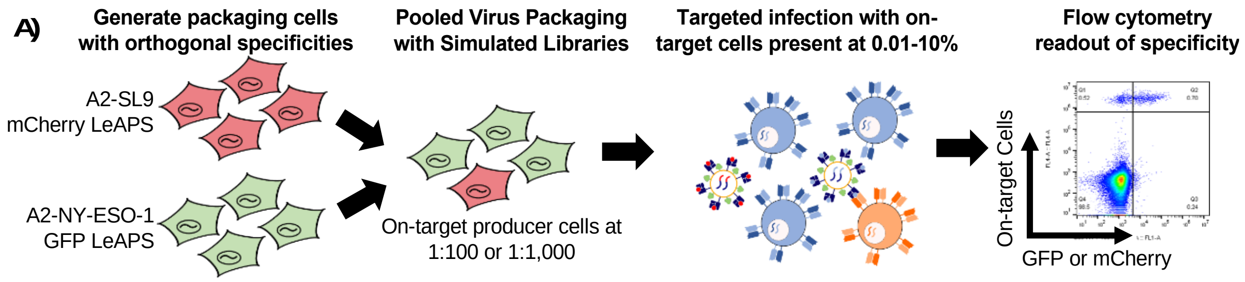


**Supplementary Figure 3 | Schematic diagram of LeAPS proof of concept experiments**

Packaging cells transduced with (i) SL9 presented on HLA-A2 and LeAPS-mCherry and (ii) NY-ESO-1 presented on HLA-A2 and LeAPS-GFP were mixed at either 1:100 or 1:1000 for each to represent libraries containing 100 constructs and 1,000 constructs, respectively. These mixtures were then used as packaging lines, transfected with helper plasmids to produce viruses, and used to transduce mixtures of cell tracking dye-labeled J76 cells expressing either 868 TCR or 1G4wt TCR at defined ratios.

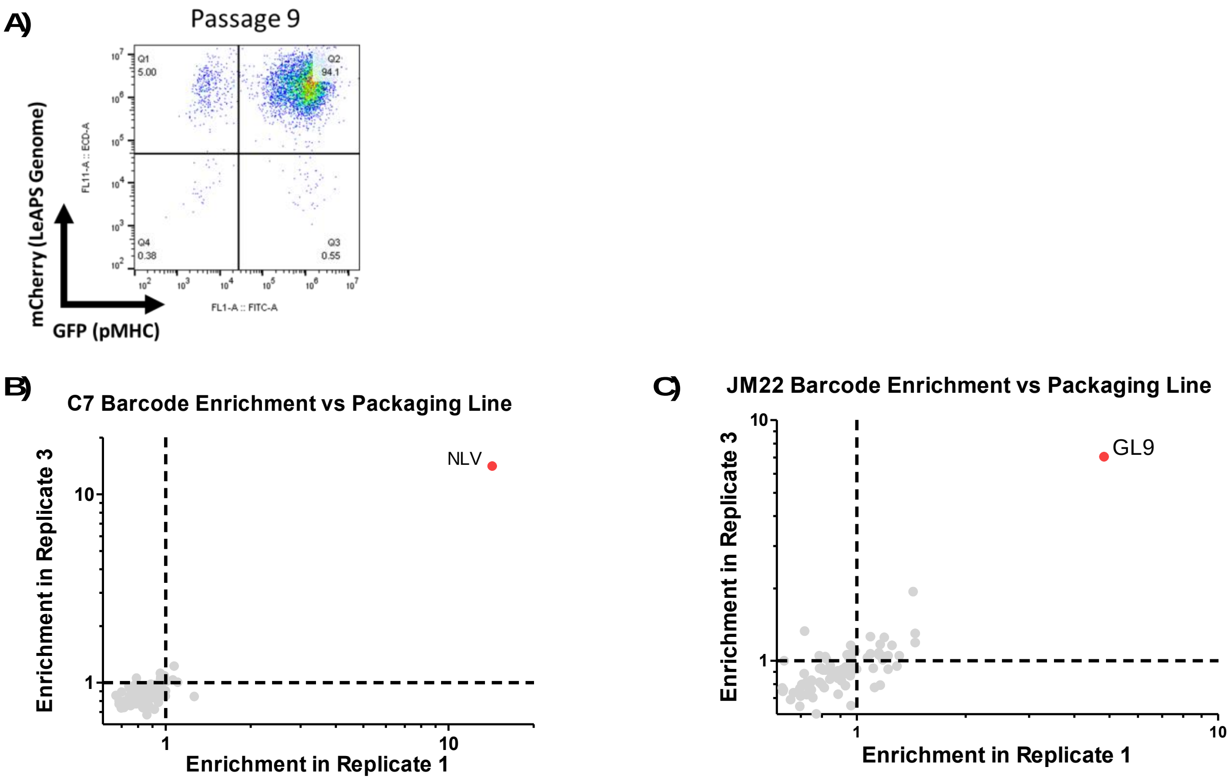


**Supplementary Figure 4 | Quality control of LeAPS packaging line and reproducibility of infection**

**A,** LeAPS-barcode expression (mCherry) and pMHC expression (GFP) of the RAPTR 96-member packaging line after 9-passages post-sorting. **B,** Correlation between infection replicates of the C7 TCR cell line. **C,** Correlation between infection replicates of the JM22 TCR cell line.

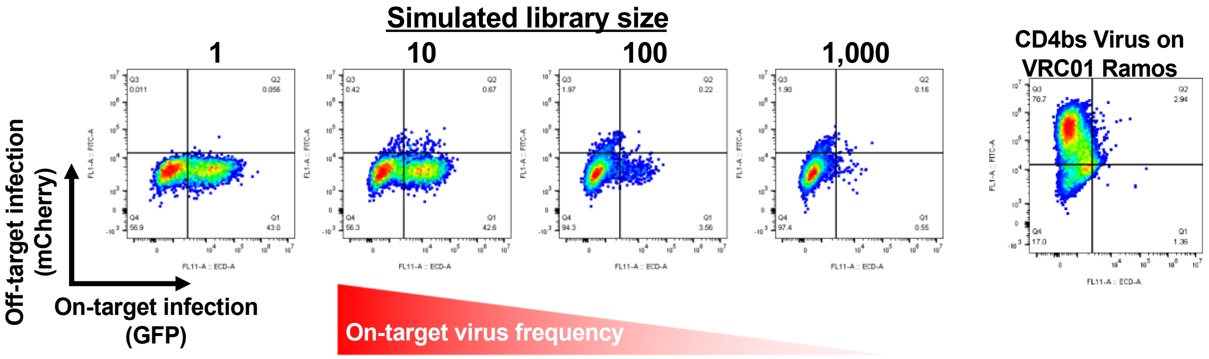


**Supplementary Figure 5 | Signal to noise in virus mixtures containing SARS-CoV-2 full spike and HIV env**

Infection of CR3022 BCR-expressing Ramos cells with mixtures of SARS-CoV-2 prefusion stabilized spike (on-target, expressing mCherry) and HIV env (off-target, expressing GFP) viruses; variant library size indicates the ratio of off-target to on-target virus present. Infection of VRC01-expressing Ramos cells with HIV env hybrid pseudotyped viruses (right) demonstrates that the viral particles remain functional.

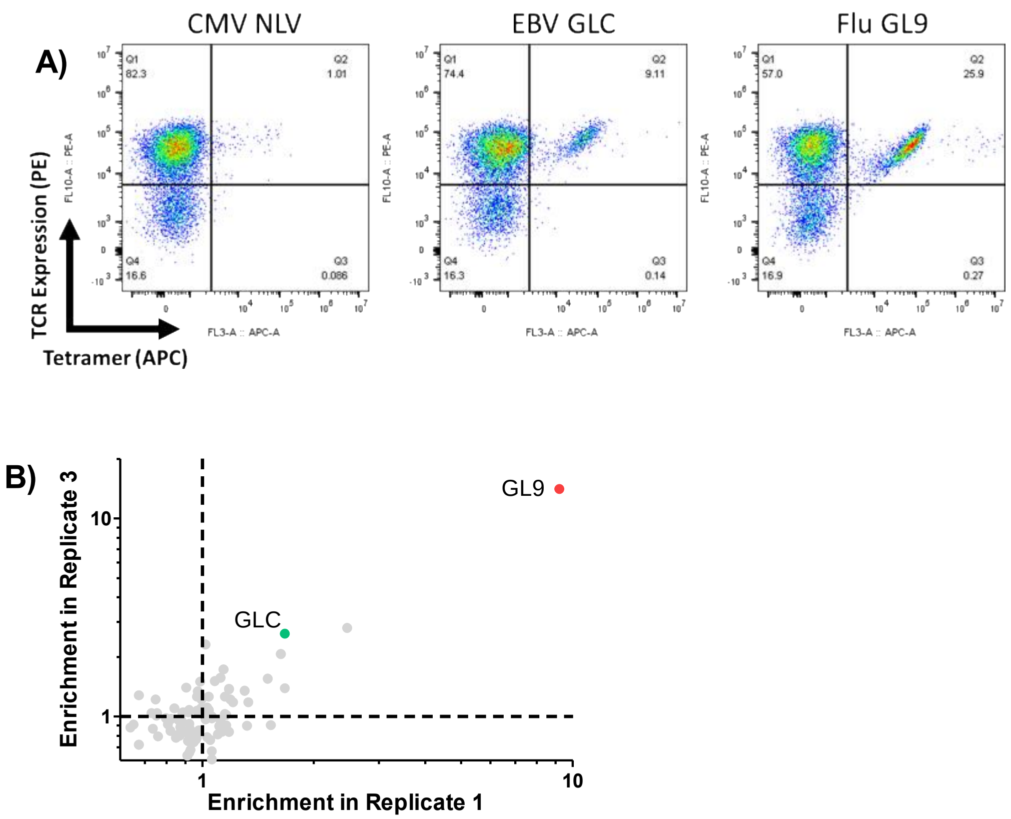


**Supplementary Figure 6 | Quality control and reproducibility for the library vs library experiment**

**A)** Tetramer staining for each of the pooled specificities used for enrichment. **B)** Correlation between infection replicates in bulk sequencing data for the library vs library experiment

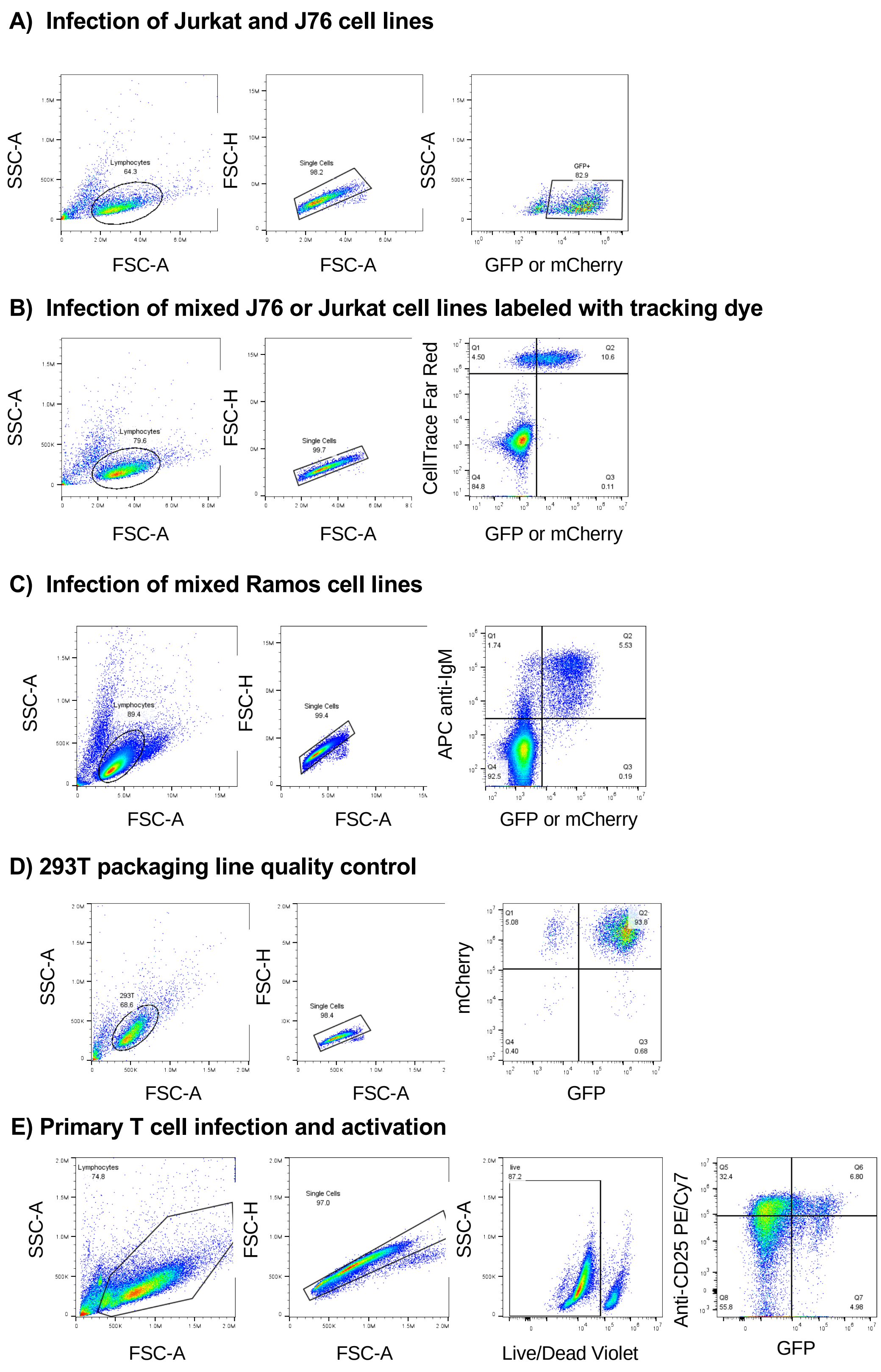


**Supplementary Figure 7 | Gating strategies used throughout manuscript**

**Supplementary Table 1 | List of peptides, barcodes, and sources used in 96-member T cell RAPTR library**

| Number | Barcode | Sequence | Parent Organism |
|---|---|---|---|
| 1 | ATTCTCCC | LLYANSAHAL | Human adenovirus 5 |
| 2 | TGCGACTC | VLAWTRAFV | Human adenovirus 5 |
| 3 | ATAATCTA | TLLYVLFEV | Human adenovirus 5 |
| 4 | AGAGAACC | RILGVLVHL | HSV-1 |
| 5 | ACACACCG | RLTGYPAGI | HSV-1 |
| 6 | ACGTTAGT | ALMLRLLRI | HSV-1 |
| 7 | CTGTTAAT | FLGGHVAVA | HSV-1 |
| 8 | TCGATCAA | TLRGLFFSV | HSV-1 |
| 9 | GTCCTGCC | SVYPYDEFV | HSV-1 |
| 10 | TGTCATCT | FLGDDPSPA | HSV-1 |
| 11 | TTGCACGG | ALLDRDCRV | HSV-1 |
| 12 | TTAAAAGC | RLLGFADTV | HSV-1 |
| 13 | CCATAAGC | ALHTALATV | HSV-1 |
| 14 | CTTCCCAT | MMLRDRWSL | HSV-1 |
| 15 | ACTCACTT | FIPQYLSAV | HSV-1 |
| 16 | CCCAACTC | FLGAGALAV | HSV-1 |
| 17 | AGACAGAT | SQLAHLVYV | HSV-1 |
| 18 | TGGGAGAG | ALMGAVTSL | HSV-1 |
| 19 | CCCTGCGC | RLNELLAYV | HSV-1 |
| 20 | GACCTAAT | TLLELVVSV | HSV-1 |
| 21 | CCCCGCTC | FLADAVVRL | HSV-1 |
| 22 | TACTATTA | FLIAYQPLL | HSV-1 |
| 23 | ACAATGTC | FLWEDQTLL | HSV-1 |

| | | | |
|---|---|---|---|
| 24 | TTCCCGCG | ILIEGIFFA | HSV-1 |
| 25 | ACGTATTA | LLTTPKFTV | HSV-1 |
| 26 | ACAGATTT | FLTCTDRSV | HSV-1 |
| 27 | TATCTGTC | GIFEDRAPV | HSV-1 |
| 28 | CTTCAGTT | FVLATGDFV | HSV-1 |
| 29 | AAACATGC | ALSALLTKL | HSV-1 |
| 30 | CTACACCT | TMYYKDVTV | HSV-1 |
| 31 | AGTATCCG | TMLEDHEFV | HSV-1 |
| 32 | ATCCATAT | ALLGLTLGV | HSV-1 |
| 33 | GTGTACCT | RMLGDVMAV | HSV-1 |
| 34 | GCGGTGAA | YLANGGFLI | HSV-1 |
| 35 | CCCGACGA | GLADTVVAC | HSV-2 |
| 36 | ACCCTTGG | FLVDAIVRVA | HSV-2 |
| 37 | AGGCGCAA | GFLIAYQPLL | HSV-2 |
| 38 | GGCCTGTC | FLVDAIVRV | HSV-2 |
| 39 | GTTAATCA | SLPRSRTPI | VZV |
| 40 | ACTCAAAG | ALWALPHAA | VZV |
| 41 | AATCCTTA | ILIEGIFFV | VZV |
| 42 | TAACCATA | MILIEGIFFV | VZV |
| 43 | GGGATTTT | RNLVPMVATVQ | CMV |
| 44 | GCGAGCTG | CYVLEETSVML | CMV |
| 45 | AAAAATAC | VLEETSVML | CMV |
| 46 | ATAAAACA | YILEETSVML | CMV |
| 47 | GGCAGGAT | NLVPMVATV | CMV |
| 48 | AGGCTGGA | FMDILTTCV | CMV |
| 49 | CCAAGCCC | LITGRLAAL | Human coronavirus 229E |
| 50 | ATAACGTC | LLLNCLWSV | Human coronavirus 229E |

| | | | |
|---|---|---|---|
| 51 | ACAGGAGA | KLWHYCSTL | Human coronavirus OC43 |
| 52 | GGTTCTAT | RFIAQLLLL | EBV |
| 53 | AATTCCCG | GLGTLGAAL | EBV |
| 54 | CTTATAAA | TLTSYWRRV | EBV |
| 55 | GCGGACAT | KLGPGEEQV | EBV |
| 56 | CGACAGAA | ALLVLYSFA | EBV |
| 57 | GGATGGGG | CLGGLLTMV | EBV |
| 58 | ATCCGTAG | LLDFVRFMGV | EBV |
| 59 | CGATGTAG | FLGERVTLT | EBV |
| 60 | TGAGTCTG | VLQWASLAV | EBV |
| 61 | GACTTGTT | TLDTKPLSV | EBV |
| 62 | TGCTTTTT | SLVIVTTFV | EBV |
| 63 | CAAATCGA | TLFIGSHVV | EBV |
| 64 | GCTGCTTA | LMIIPLINV | EBV |
| 65 | GAGTGCAC | FMVFLQTHI | EBV |
| 66 | GAGATGCA | LLWTLVVLL | EBV |
| 67 | TGTTCCGA | LLSAWILTA | EBV |
| 68 | CAACAGCG | YLQQNWWTL | EBV |
| 69 | GAATGTAT | FLYALALLL | EBV |
| 70 | AACAATTG | WQWEHIPPA | EBV |
| 71 | TTAATGAA | YLLEMLWRL | EBV |
| 72 | GCATCGGC | LLIEGIFFI | EBV |
| 73 | CTTTAAAT | IGLITVLFL | HHV8 |
| 74 | AGTCAAAA | LVLILYLCV | HHV8 |
| 75 | ACAGGGGT | LLNGWRWRL | HHV8 |
| 76 | CGCTCATT | GLCTLVAML | EBV |
| 77 | GAGCCATT | TLDYKPLSV | EBV |

| | | | |
|---|---|---|---|
| 78 | AGCTTAAT | FLDKGTYTL | EBV |
| 79 | GCCAAAAA | YVLDHLIVV | EBV |
| 80 | CACACACT | VLFGLLCLL | EBV |
| 81 | TTAATCAA | GILGFVFTL | Influenza A virus |
| 82 | TGAACGCC | NMLSTVLGV | Influenza A virus |
| 83 | CTTGATGG | RLYQNPTTYI | Influenza A virus |
| 84 | GACTGCCA | KLYQNPTTYI | Influenza A virus |
| 85 | ATTTCCGG | CVNGSCFTV | Influenza A virus |
| 86 | GGTTGCGT | GILGFVFTLT | Influenza A virus |
| 87 | ACTGCATT | KLWESPQEI | Measles morbillivirus |
| 88 | AAAGAGTA | ILPGQDLQYV | Measles morbillivirus |
| 89 | GGACCAAA | SMYRVFEVGV | Measles morbillivirus |
| 90 | CGCGTAAG | FMYMSLLGV | Measles morbillivirus |
| 91 | TGAAGAGG | GMNVANHFL | Mumps rubulavirus |
| 92 | TGGGTTGA | ALDQTDIRV | Mumps rubulavirus |
| 93 | TACAATTG | LLDSSTTRV | Mumps rubulavirus |
| 94 | GCATCCCG | VMNQVIHGV | Mumps rubulavirus |
| 95 | GCTTAGAC | GLMEGQIVSV | Mumps rubulavirus |
| 96 | TCAAGTGC | KLIAGVNYI | Mumps rubulavirus |

**Supplementary Table 2 | List of viral surface glycoproteins and immunogens, barcodes, and modifications used in B cell RAPTR library**

| Number | Name | Barcode | Modifications |
|---|---|---|---|
| 1 | SARS-CoV-2 Wuhan Strain | ATTCTCCC | Prefusion stabilized (2P) |
| 2 | D614G | TGCGACTC | Prefusion stabilized (2P) |
| 3 | B.1.1.7 | ATAATCTA | Prefusion stabilized (2P) |
| 4 | B.1.1.298 | AGAGAACC | Prefusion stabilized (2P) |
| 5 | B.1.429 | ACACACCG | Prefusion stabilized (2P) |
| 6 | P.2 | ACGTTAGT | Prefusion stabilized (2P) |
| 7 | P.1 | CTGTTAAT | Prefusion stabilized (2P) |
| 8 | B.1.351 v1 | TCGATCAA | Prefusion stabilized (2P) |
| 9 | B.1.351 v2 | GTCCTGCC | Prefusion stabilized (2P) |
| 10 | B.1.351 v3 | TGTCATCT | Prefusion stabilized (2P) |
| 11 | SARS-CoV-1 | TTGCACGG | Prefusion stabilized (2P) |
| 12 | WIV1-CoV | TTAAAAGC | Prefusion stabilized (2P) |
| 13 | OC43 | CCATAAGC | Prefusion stabilized (2P) |
| 14 | 229E | CTTCCCAT | Prefusion stabilized (2P) |
| 15 | HKU1 | ACTCACTT | Prefusion stabilized (2P) |
| 16 | NL63 | CCCAACTC | Prefusion stabilized (2P) |
| 17 | MERS | AGACAGAT | Prefusion stabilized (2P) |
| 18 | B.1.160 | TGGGAGAG | Prefusion stabilized (2P) |
| 19 | B.1.525 | CCCTGCGC | Prefusion stabilized (2P) |
| 20 | B.1.526 | GACCTAAT | Prefusion stabilized (2P) |
| 21 | B.1.526.1 | CCCCGCTC | Prefusion stabilized (2P) |
| 22 | B.1.526.2 | TACTATTA | Prefusion stabilized (2P) |
| 23 | B.1.258 | ACAATGTC | Prefusion stabilized (2P) |
| 24 | B.1.221 | TTCCCGCG | Prefusion stabilized (2P) |

| | | | |
|---|---|---|---|
| 25 | B.1.1.277 | ACGTATTA | Prefusion stabilized (2P) |
| 26 | B.1.1.302 | ACAGATTT | Prefusion stabilized (2P) |
| 27 | B.1.367 | TATCTGTC | Prefusion stabilized (2P) |
| 28 | B.1.617 | CTTCAGTT | Prefusion stabilized (2P) |
| 29 | MHV | AAACATGC | Prefusion stabilized (2P) |
| 30 | SARS 2 E | AGTATCCG | |
| 31 | SARS 2 M | ATCCATAT | |
| 32 | H1 | GTGTACCT | Y98F to abrogate sialic acid binding |
| 33 | H3 | GCGGTGAA | Y98F to abrogate sialic acid binding |
| 34 | N1 | CCCGACGA | Y406D to reduce enzymatic activity |
| 35 | N2 | ACCCTTGG | Y406D to reduce enzymatic activity |
| 36 | Measles H | AGGCGCAA | Intracellular domain truncated 18aa |
| 37 | Measles F | GGCCTGTC | Intracellular domain truncated 30aa |
| 38 | HIV SOSIP (PDGFR TM/Stalk) | GTTAATCA | |
| 39 | CMV gB | ACTCAAAG | |
| 40 | EBV gB | AATCCTTA | |
| 41 | RSV F DS-Cav1 (PDGFR TM/Stalk) | TAACCATA | Prefusion stabilized |
| 42 | Nipah Virus G | GGGATTTT | Intracellular domain truncated 34aa |
| 43 | Dengue Virus Type 2 E Monomer | GCGAGCTG | Residues 2-396, On PDGFR TM + stalk |
| 44 | Dengue Virus Type 2 Dimer | AAAAATAC | Residues 2-394, linked with $(G_4S)_3$, on PDGFR TM + Stalk |

**Supplementary Methods**

1) Add custom primers into RT Mastermix, then perform droplet capture with the 10x Genomics Chromium Single Cell 5' Chemistry Kits.

Prior to rt, add 0.5uL of 10uM Barcode RT primer directly into the 68.3uL of Mastermix for each lane.

2) cDNA amplification

Each of the custom primers has an adapter identical to the adapter sequence on the Poly-dT RT Primer (10x Genomics, PN-2000007). This adapter servers as a primer binding site for the NonPoly(dT) primer (10x Genomics, PN-220106) during cDNA amplification, and thus allows amplification of reverse transcribed TCR and barcode products. Therefore, the cDNA amplification can be performed according to the 10x kit instructions.

We expect a 1,000 bp amplicon on a BioAnalyzer from the custom primer.

3) PCR amplification of each transcript and sample indexing

Prepare TCR amplicons according to the standard V(D)J kit instructions.

<u>For the mCherry Barcode constructs:</u>

First PCR to add i7 handle for each sample (do 4 parallel reactions for each sample and then pool):

| **Reagent** | **Volume for 1 Reaction** | **Volume for 4.25 Reactions** |
|---|---|---|
| 10uM mChBC_ui7_r | 1 | 4.25 |
| 10uM P5 | 1 | 4.25 |
| Amplified cDNA: 10ng | Depends on DNA conc | Depends on DNA conc |
| 2x KAPA Hi-Fix Master Mix | 25 | 106.25 |
| Water | Depends on DNA conc | Depends on DNA conc |
| Total (uL) | 50 | 212.5 |

<u>PCR 1 conditions:</u>

| **Step** | **Time** | **Temperature** | **Cycles** |
|---|---|---|---|
| Initial Denaturation | 3 min | 95°C | 1 |
| Denature | 15 s | 98°C | 25 |
| Anneal | 15 s | 57°C | |
| Extend | 1 min | 72°C | |
| Final Elongation | 5 min | 72°C | 1 |
| Hold | Indefinitely | - | 1 |

Purification for PCR 1

Perform 0.6x left-sided SPRI cleanup (taking the elution from the beads) to recover amplified DNA and remove primers.

Expected size is ~900 bp. Refer to this as mCherryBC-i7 and carry each forward to next step.

Second PCR to add sequencing index to barcodes (do 4 parallel reactions for each sample and then pool):

| **Reagent** | **Volume for 1 Reaction** | **Volume for 4.25 Reactions** |
|---|---|---|
| 10uM BC0X (X = 1 for sample 1, 2 for sample 2) | 1 | 4.25 |
| 10uM P5 | 1 | 4.25 |
| Amplified mCherryBC-i7: 10ng | Depends on DNA conc | Depends on DNA conc |
| 2x KAPA Hi-Fix Master Mix | 25 | 106.25 |
| Water | Depends on DNA conc | Depends on DNA conc |
| Total (uL) | 50 | 212.5 |

PCR 2 conditions:

| **Step** | **Time** | **Temperature** | **Cycles** |
|---|---|---|---|
| Initial Denaturation | 3 min | 95°C | 1 |
| Denature | 15 s | 98°C | 25 |
| Anneal | 15 s | 57°C | |
| Extend | 1 min | 72°C | |
| Final Elongation | 5 min | 72°C | 1 |
| Hold | Indefinitely | - | 1 |

Purification for PCR 2 and Sequencing Prep

Perform 0.6x left-sided SPRI cleanup (taking the elution from the beads) to recover amplified DNA and remove primers. Expected size is ~920bp.

This is now indexed DNA ready for QC and sequencing on the MiSeq, with read length 150nt.

4) Sequencing
   TCRs from all V(D)J preps should be pooled and run on a MiSeq or NextSeq.

   The mCherry Amplicon samples should be analyzed on 2 MiSeq runs (4 pooled samples each). For the mCherryBC amplicon: 150nt (75+75) paired end.

Sequencing primers are Solexa i5 and i7. Primer sequences are below for verification.

| Name | Full Primer Sequence (5' to 3') | Barcode Sequence |
|---|---|---|
| **RT Spike-in** | AAGCAGTGGTATCAACGCAGAGTACGAGGAGAAAATGAAAGCCATACGGGAAGC | - |
| **mCh_BC_ui7_r** | GCTGAACCGCTCTTCCGATCTNNNNNNNNCAGAGGTTGATTACCGATAAGCTTGATATCG | - |
| **BC01** | CAAGCAGAAGACGGCATACGAGATATCACGCGGTCTCGGCATTCCTGCTGAACCGCTCTTCCGATCT | CGTGAT |
| **BC02** | CAAGCAGAAGACGGCATACGAGATCGATGTCGGTCTCGGCATTCCTGCTGAACCGCTCTTCCGATCT | ACATCG |
| **BC03** | CAAGCAGAAGACGGCATACGAGATTTAGGCCGGTCTCGGCATTCCTGCTGAACCGCTCTTCCGATCT | GCCTAA |
| **BC04** | CAAGCAGAAGACGGCATACGAGATTGACCACGGTCTCGGCATTCCTGCTGAACCGCTCTTCCGATCT | TGGTCA |
| **BC05** | CAAGCAGAAGACGGCATACGAGATACAGTGCGGTCTCGGCATTCCTGCTGAACCGCTCTTCCGATCT | CACTGT |
| **BC06** | CAAGCAGAAGACGGCATACGAGATGCCAATCGGTCTCGGCATTCCTGCTGAACCGCTCTTCCGATCT | ATTGGC |
| **BC07** | CAAGCAGAAGACGGCATACGAGATCAGATCCGGTCTCGGCATTCCTGCTGAACCGCTCTTCCGATCT | GATCTG |
| **BC08** | CAAGCAGAAGACGGCATACGAGATACTTGACGGTCTCGGCATTCCTGCTGAACCGCTCTTCCGATCT | TCAAGT |
| **P5 (fwd)** | AATGATACGGCGACCACCGAGATCTACACACACTCTTTCCCTACACGACGCTCTTCCGATCT | - |
